# Shaky Scaffolding: Age Differences In Cerebellar Activation Revealed Through Activation Likelihood Estimation Meta-Analysis

**DOI:** 10.1101/716530

**Authors:** Jessica A. Bernard, An D. Nguyen, Hanna K. Hausman, Ted Maldonado, Hannah K. Ballard, T. Bryan Jackson, Sydney M. Eakin, Yana Lokshina, James R.M. Goen

**Author notes:** **Corresponding Author** Jessica A. Bernard, PhD, Department of Psychological and Brain Sciences, Texas A&M University. **Data Availability:** All materials associated with the analysis in the form of text files of foci are freely available for download at https://osf.io/gx5jw/. **Ethics Statement:** This investigation used published, anonymous data and as such was not subject to ethics review.

## Abstract

Cognitive neuroscience research has provided foundational insights into aging, but has focused primarily on the cerebral cortex. However, the cerebellum is subject to the effects of aging. Given the importance of this structure in the performance of motor and cognitive tasks, cerebellar differences stand to provide critical insights into age differences in behavior. But, our understanding of cerebellar functional activation in aging is limited. Thus, we completed a meta-analysis of neuroimaging studies across task domains. Unlike in the cortex where an increase in bilateral activation is seen during cognitive task performance with advanced age, there is less overlap in cerebellar activation across tasks in older adults relative to young. Conversely, we see an increase in activation overlap in older adults during motor tasks. We propose that this is due to inputs for comparator processing in the context of control theory (cortical and spinal) that may be differentially impacted in aging. These findings advance our understanding of the aging mind and brain.

## Introduction

Advanced age is accompanied by differences in both cognitive (e.g., Park, Polk, Mikels, Taylor, & Marshuetz, 2001) and motor behavior (reviewed in Seidler et al., 2010). The impact of these differences on quality of life, well-being, independence, and rehabilitation is large. For example, cognitive complaints and deficits in memory, even in cognitively normal older adults (OA), have broad impacts on quality of life across social and cognitive domains (Parikh, Troyer, Maione, & Murphy, 2016). Motor systems differences include those in learning abilities (e.g., Anguera, Reuter-Lorenz, Willingham, & Seidler, 2011) and fall risk. Falls are a major cause of disability in OA, and postural control is associated with cognitive performance (Huxhold, Li, Schmiedek, & Lindenberger, 2006). Characterizing the neural underpinnings of age-related cognitive and motor behavioral differences is critical for both our basic understanding of the aging process and for the elucidation of new remediation targets to improve quality of life for OA.

To this end, the field of the cognitive neuroscience of aging has greatly advanced our understanding of how brain changes and differences in OA impact behavior. Broadly, we know that in advanced age, the brain is smaller (e.g., Walhovd et al., 2011), there are differences in functional networks at rest (Andrews-Hanna et al., 2007; Langan et al., 2010), and differences in brain activation patterns during task performance (Naccarato et al., 2006; Reuter-Lorenz, Stanczak, & Miller, 1999; Seidler et al., 2010). Patterns of bilateral activation in OA are commonly seen in cases where young adults (YA) would typically only activate one hemisphere (Cabeza, 2002; Reuter-Lorenz et al., 1999). This bilateral activation, particularly in the prefrontal cortex (PFC), has been suggested to be compensatory (Cabeza, 2002; Reuter-Lorenz & Cappell, 2008) to help maintain performance in advanced age.

While investigating cortical differences in OA has critically informed our understanding of age-related performance differences, cerebellar contributions to performance have been relatively understudied. There is a growing literature which demonstrates age differences in cerebellar volume (J. A. Bernard & Seidler, 2013b; Koppelmans, Young, & Sarah, 2017; Miller et al., 2013; Raz et al., 2005) and connectivity with the cortex (Bernard et al., 2013). As our understanding of the functional contributions of the cerebellum has grown, we know now that it contributes to both motor and cognitive task performance (e.g., Balsters, Whelan, Robertson, & Ramnani, 2013; Chen & Desmond, 2005; Stoodley & Schmahmann, 2009). With these wide-ranging behavioral contributions, understanding how the cerebellum may contribute to performance in OA is of great interest and importance. To date, work in this area has demonstrated that differences in both volume and connectivity of the cerebellum are functionally relevant for both motor and cognitive performance in OA (Bernard & Seidler, 2013; Bernard et al., 2013; Miller et al., 2013).

Recently, we suggested that differences in connectivity with the cortex and smaller lobular volume in the cerebellum in OA contribute to performance differences in aging due to degraded internal models of behavior (Bernard & Seidler, 2014). Theories of cerebellar function have suggested that the structure acts on copies of commands for behavior and compares the outcomes of a given command with what is expected based on that initial command (Ramnani, 2006, 2014) Ultimately, internal models of a particular movement or thought process are formed (e.g., Balsters et al., 2013; Imamizu et al., 2000) that allow for greater automaticity. However, due to degraded cerebello-cortical connectivity and the smaller cerebellar volume in OA, the inputs to this structure may be negatively impacted, resulting in degraded internal model processing and, in turn, performance deficits (Bernard & Seidler, 2014). Further, cerebellar function may provide important scaffolding for behavior in OA (Filip, Gallea, Lehéricy, Lungu, & Bareš, 2019). Indeed, it may be the case that the bilateral processing seen in the cortex in OA is to compensate for a lessened ability to rely upon cerebellar resources in advanced age. That is, the brain is less able to offload processing and take advantage of more automatic processing via existing internal models, resulting in a greater need for cortical resources. However, there has been limited work investigating the functional activation patterns of the cerebellum in advanced age, which would provide important insight into this hypothesis. Investigating cerebellar functional activation in OA stands to advance our understanding of the neural underpinnings of behavioral differences and changes in advanced age, providing a more complete perspective on the aging mind and brain.

To better understand the functional engagement of the cerebellum in OA we conducted a meta-analysis of the functional brain imaging literature in OA and YA. We tested two competing hypotheses. Based on our prior work suggesting degraded inputs to the cerebellum and internal model processing in OA (Bernard & Seidler, 2014), decreased convergence of cerebellar activation in OA relative to YA would be expected. That is, we would expect to see less consistent overlap in foci of activation across studies. However, the cortical literature consistently demonstrates an increase in bilateral activation in OA during task performance (e.g., Cabeza, 2002; Reuter-Lorenz et al., 1999). Thus, alternatively, the same pattern may be present in the cerebellum if similar compensatory processes are recruited during task processing. As such, we would expect to see a convergence of foci across studies in both cerebellar hemispheres in OA, while in YA convergence would be limited to one hemisphere (consistent with lateralized findings from prior investigations of the cerebellar functional topography; e.g., E et al., 2012; Stoodley et al., 2012; Stoodley and Schmahmann, 2009b).

## Method

### Literature Search and Inclusion Criteria

All materials associated with the analysis in the form of text files of foci (for more details see below) are freely available for download at https://osf.io/gx5jw/. To identify papers, we completed two separate and sequential literature searches completed using PubMed (http://www.ncbi.nlm.nih.gov/pubmed). The first search used the search term: “cerebell* AND imaging” with the limits “Humans” and “English.” Additionally, we included the limit “Adult 65+” to target the OA literature. This resulted in 3,913 articles.

Articles that focused on structural or morphometric analyses, region of interest analysis, and functional connectivity, as well as those that did not report coordinates in the cerebellum, did not report coordinates in standard spaces (Montreal Neurological Institute (MNI) or Talairach), and did not have independent groups contrast analysis were excluded. This is consistent with the exclusion criteria used in recent meta-analyses from our group, and others (Bernard & Mittal, 2015; Bernard, Russell, Newberry, Goen, & Mittal, 2017; Bernard & Seidler, 2013a; E et al., 2012; Stoodley & Schmahmann, 2009). After completion of this search and exclusion of papers based on the aforementioned exclusion criteria, we were left with a very small sample of studies (43 studies) and foci on which to complete our analyses (see Figure 1). However, this was limited, at least in part, to the inclusion of the “cerebell*” term in our initial search, as this term may not be in the keywords or abstracts of papers indexed in PubMed. As such, we completed a second search using the terms “aging AND brain imaging” with the same limits as above, which returned 5,982 results as of August 6, 2018. All inclusion/exclusion criteria were identical those for search 1. This second search yielded an additional 73 studies for inclusion.

**Figure 1.**
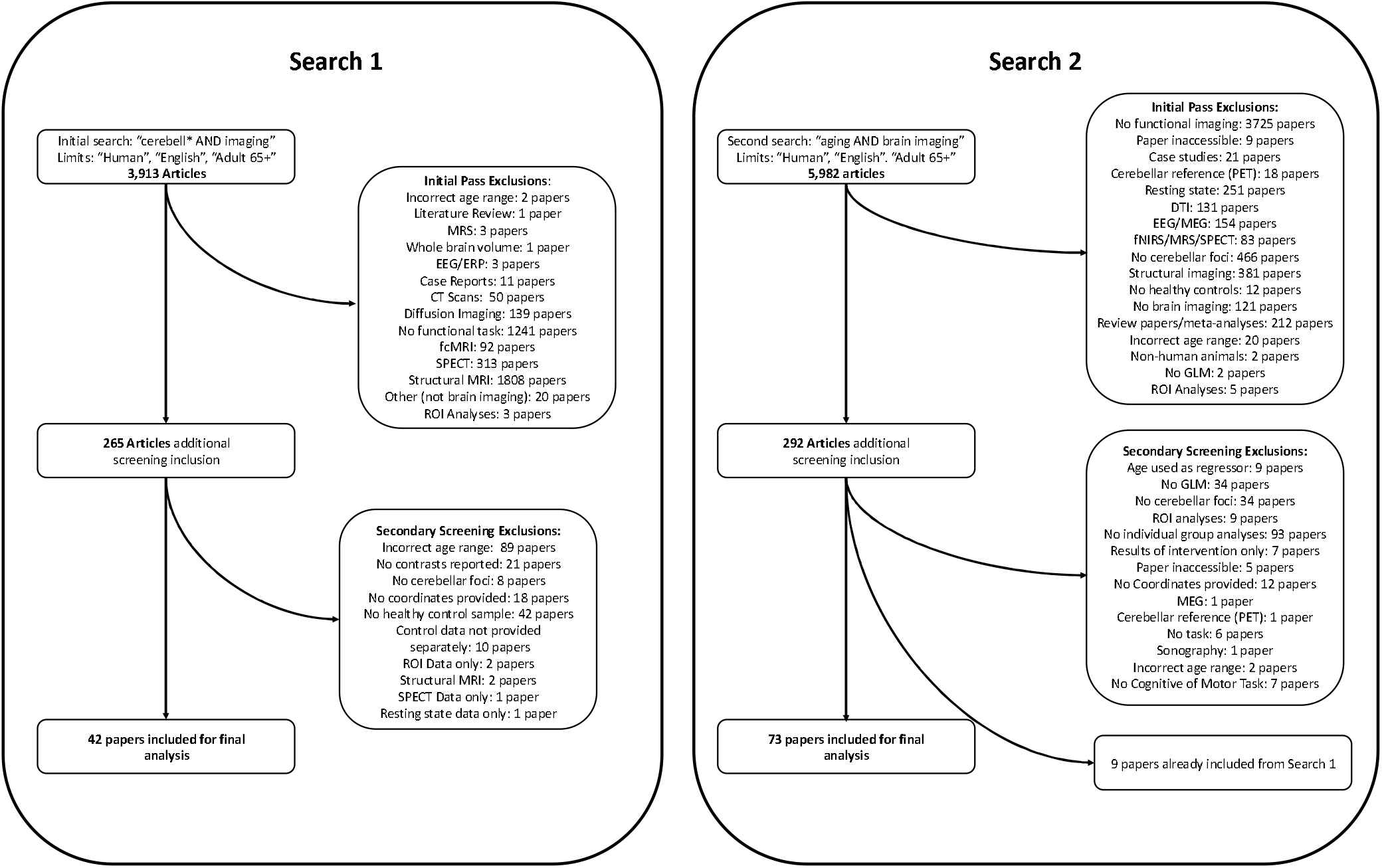
Flow chart describing the two search processes. In both cases, an initial overview of papers was conducted to eliminate initial obvious exclusions. Secondary screening was conducted while foci were pulled from papers for analysis, though additional exclusions occurred at this stage as well. A full list of included papers can be found in Tables 1-5, organized by task domain. 115 studies were included based off of our two literature searches. However, additional studies from prior meta-analyses of cerebellar function were also added to our sample, bringing the total number of included studies to 175.

As cerebellar engagement in both motor and cognitive tasks was of interest, we included studies in the following task domains: motor function, working memory, language, and “other cognitive tasks.” Notably, this last category primarily included executive function tasks (such as the Stroop task, tower of London task, etc.), though several tasks assessing spatial processing were also included here. Studies from E and colleagues (2012) categorized as executive function were put into this category. Categorical determination was made to be consistent with the task domains used by both E and colleagues (2012) and Stoodley and Schmahmann (2009), with the exception of long-term memory as it had not been previously included in past meta-analyses. Tasks included in this category included a memory component with a delay in recall, typically on the order of several minutes. These task domains were chosen for several key reasons. First, we aimed to parallel prior meta-analyses looking at cerebellar function to compare the functional topography in OA to what is known about this topography in YA. Second, these are domains where there are known age differences in performance and as such are of great interest in the study of motor and cognitive aging. Tables 1-5 includes a complete listing of the studies included in our meta-analysis divided by task domain, along with the average age and/or age range of participants when available. This also provides information about the brain imaging modality (PET or fMRI), scanner field strength where applicable, and the number of foci from a given study for each age group.

**Table 1.**
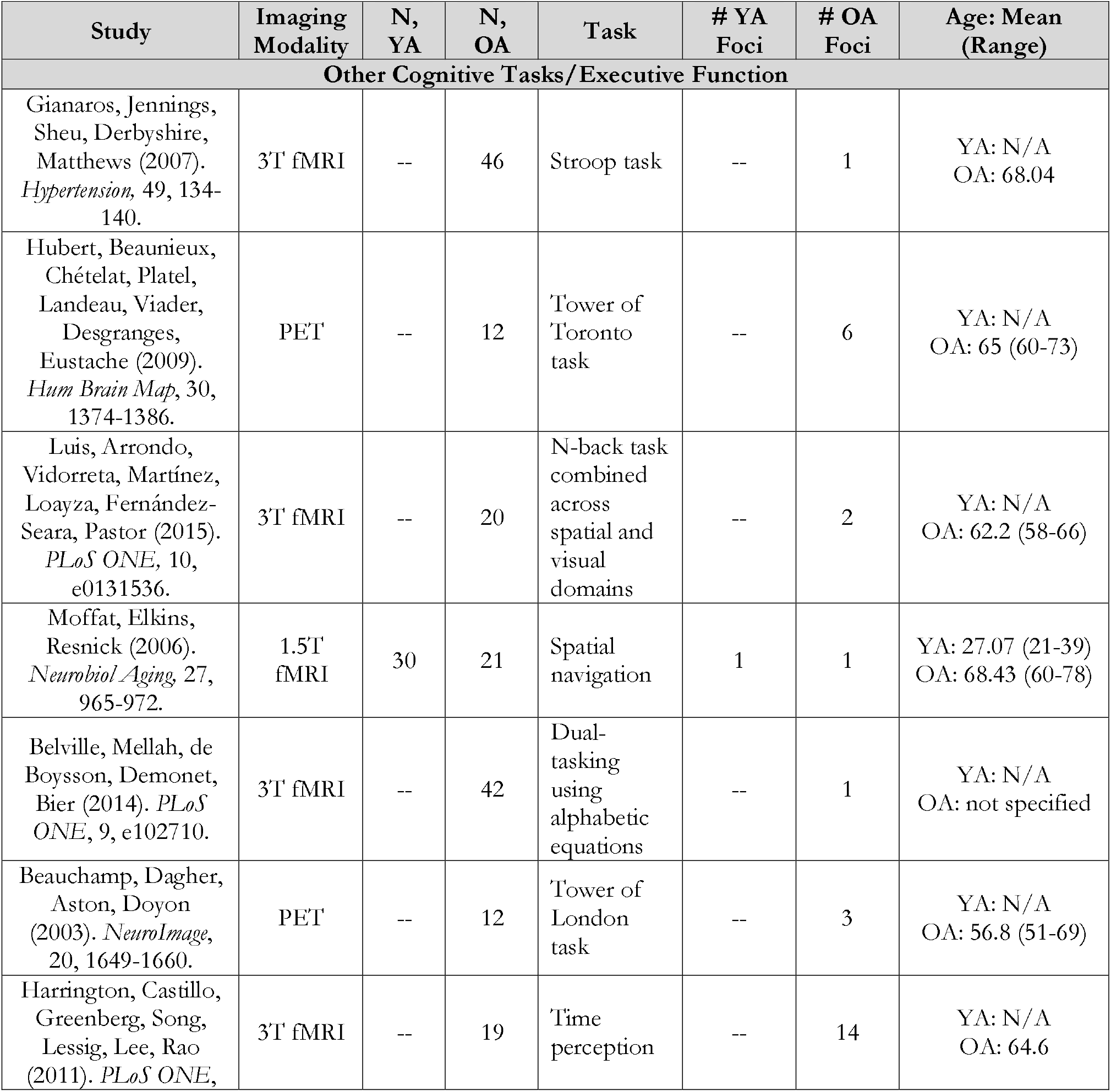
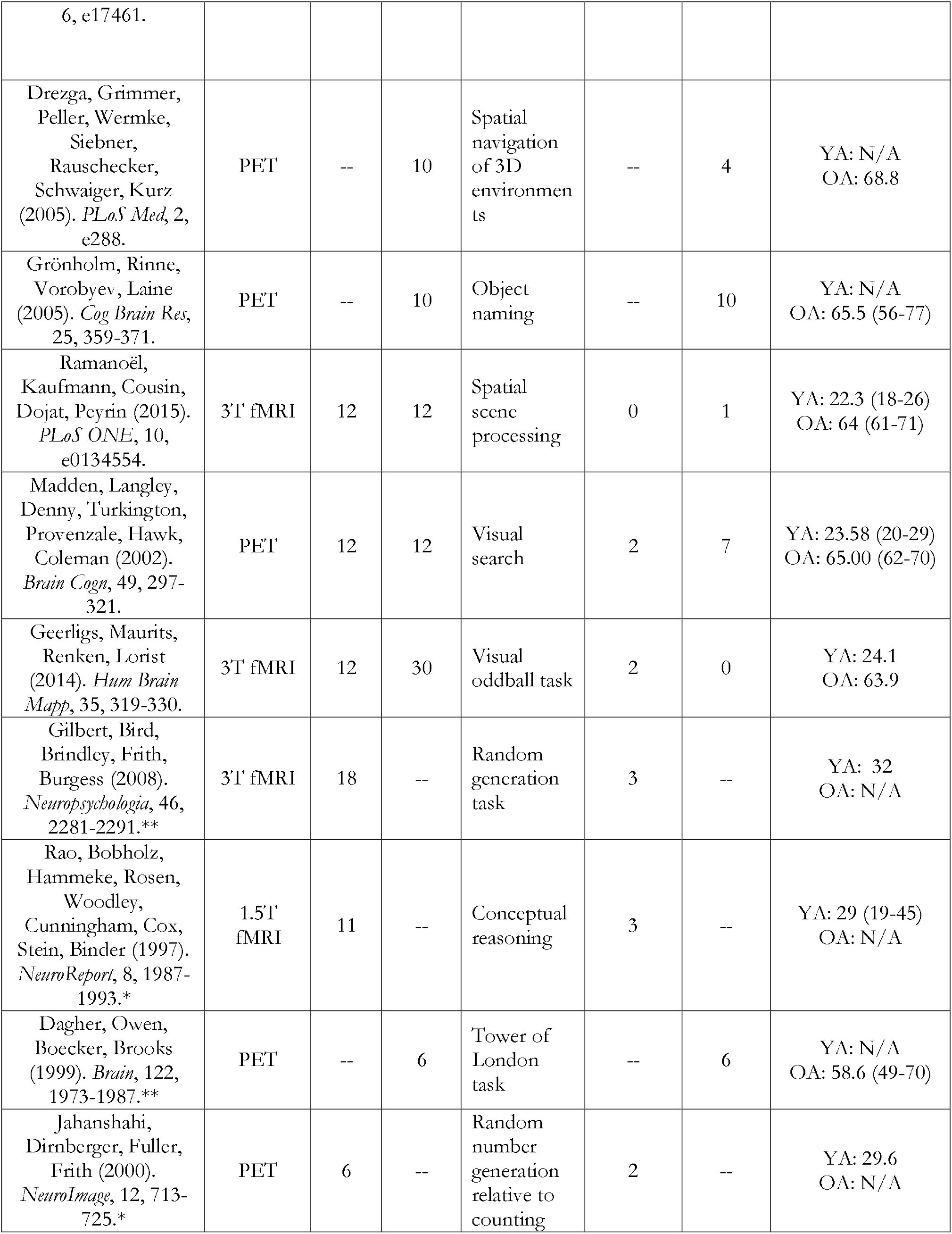
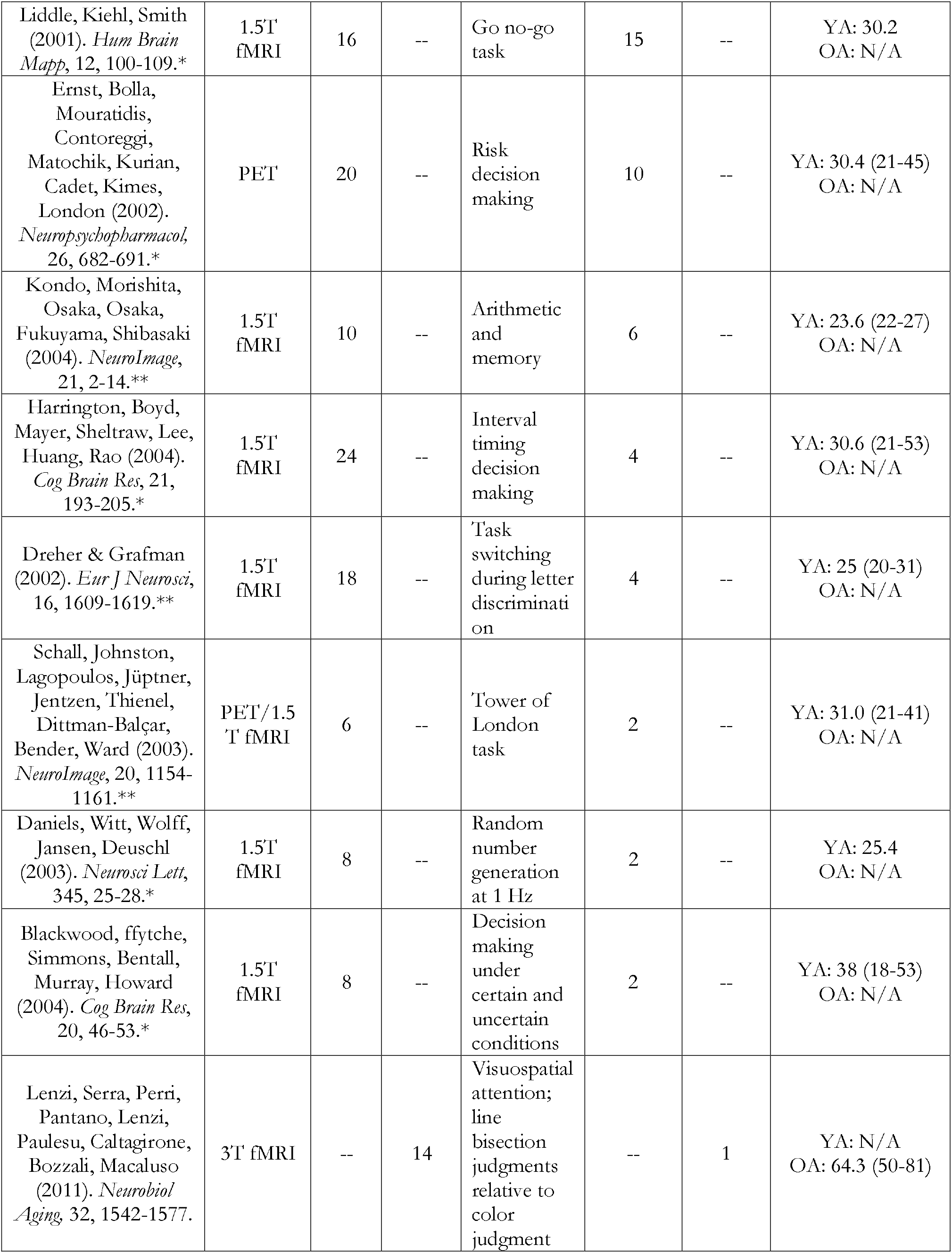
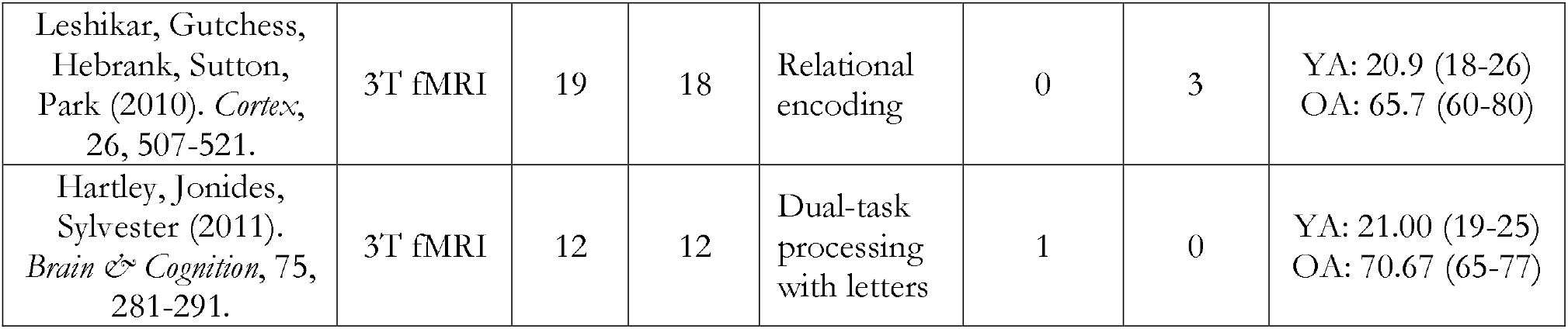
Included studies in the “Other Cognitive/Executive Function” category. Notably, while many of the included studies had both YA and OA, in some instances OA data came from clinical studies wherein the OA served as a control group. Further, additional YA data came from studies included in prior meta-analyses. ^#^Studies included only as part of Stoodley & Schmahmann, 2009. *Studies included as part of Stoodley & Schmahmann, 2009 and E et al., 2014. **Studies only included in E et al., 2014. “—” denotes studies where group a particular age group was not included and as such no coordinates are possible. Cases where there were no cerebellar coordinates are indicated by a 0 in the appropriate foci column. Mean age is provided in years, and the range is also provided when available. N/A: not applicable

**Table 2.** Included studies of language tasks. Notably, while many of the included studies had both YA and OA, in some instances OA data came from clinical studies wherein the OA served as a control group. Further, additional YA data came from studies included in prior meta-analyses. ^#^Studies included only as part of Stoodley & Schmahmann, 2009. *Studies included as part of Stoodley & Schmahmann, 2009 and E et al., 2014. **Studies only included in E et al., 2014. “—” denotes studies where group a particular age group was not included and as such no coordinates are possible. Cases where there were no cerebellar coordinates are indicated by a 0 in the appropriate foci column. N/A: not applicable

| Study | Imaging Modality | N, YA | N, OA | Task | # YA Foci | # OA Foci | Age: Mean (Range) |
| --- | --- | --- | --- | --- | --- | --- | --- |
| **Language** |  |  |  |  |  |  |  |
| Rizio, Moyer, Diaz (2017). *Brain Behav*, 7, e00660. | 3T fMRI | 20 | 20 | Language interference naming | 0 | 2 | YA: 23.7 (18-31)<br>OA: 67.25 (60-79) |
| Provost, Brambati, Chapleau, Wilson (2016). *Cortex*, 84, 90-100. | 3T fMRI | 16 | 16 | Word reading | 0 | 1 | YA: 27.5 (22-33)<br>OA: 67.0 (60-75) |
| Martins, Simard, Monchi (2014). *PLoS ONE*, 9, e99710. | 3T fMRI | 14 | 14 | Lexical version of the Wisconsin card sorting task | 6 | 3 | YA: 26 (21-31)<br>OA: 63 (55-71) |
| Whatmough, Verret, Fung, Chertkow (2004). *J Cogn Neurosci*, 16, 1211-1226. | PET | -- | 15 | Semantic judgments of word pairs | -- | 4 | YA: N/A<br>OA: 74.3 (69-90) |
| Olichney, Taylor, Hillert, Chan, Salmon, Gatherwright, Iragui, Kutas (2010). *Neurobiol Aging*, 31, 1975-1990. | 1.5T fMRI | -- | 17 | Word memory and repetition | -- | 3 | YA: N/A<br>OA: 69.7 |
| Daselaar, Veltman, Rombouts, Raaijmakers, Jonker (2003). *Neurobiol Aging*, 24, 1005-1011. | 1.5T fMRI | 26 | 39 | Semantic characterization after shallow or deep encoding | 1 | 1 | YA: 32.4 (30-35)<br>OA: 66.3 (63-71) |
| Madden, Langley, Denny, Turkington, Provenzale, Hawk, Coleman (2002). *Brain Cogn*, 49, 297-321. | PET | 12 | 12 | Lexical decision task | 3 | 2 | YA: 23.58 (20-29)<br>OA: 65.00 (62-70) |
| Seki, Okada, Koeda, Sadato (2004). *Cog Brain Res*, 20, 261-272.* | 3T fMRI | 19 | -- | Vowel exchange compared to reading words and non-words | 2 | -- | YA: 23.3<br>OA: N/A |
| Rauschecker, Pringle, Watkins (2008). *Hum Brain Mapp*, 29, 1231-1242.** | 3T fMRI | 14 | -- | Listening and cover repetition of non-words | 4 | -- | YA: 23.3 (20-34)<br>OA: N/A |
| Daselaar, Veltman, Rombouts, Raaijmakers, Jonker (2005). *Neurobiol Learn Mem*, 83, 251-262. | 1.5T fMRI | 25 | 38 | Word-stem completion | 1 | 0 | YA: 32.3 (30-35)<br>OA: 66.4 (63-71) |
| Ojemann, Buckner, Akbudak, Snyder, Ollinger, McKinstry, Rosen, Petersen, Raichle, Conturo (1998). *Hum Brain Mapp*, 6, 203-215.* | PET/1.5 T fMRI | 7 | -- | Word-stem completion | 7 | -- | YA: 24 (19-28)<br>OA: N/A |
| Schlosser, Hutchinson, Joseffer, Rusinek, Saarimaki, Stevenson, Dewey, Brodie (1998). *J Neurol Neurosurg Psychiatry*, 64, 492-498.* | 1.5T fMRI | 6 | -- | Verbal fluency task | 6 | -- | YA: 23 (22-26)<br>OA: N/A |
| Lurito, Kareken, Lowe, Chen, Mathews (2000). *Hum Brain Mapp*, 10, 99-106.* | 1.5T fMRI | 5 | -- | Word generation compared to viewing non-letter symbols | 3 | -- | YA: 27<br>OA: N/A |
| Seeger, Desmond, Glover, Gabrieli (2000). *Neuropsychol*, 14, 361-369.* | 1.5T fMRI | 7 | -- | Verb generation task | 16 | -- | YA: 31<br>OA: N/A |
| Gurd, Amunts, Weiss, Zafiris, Zilles, Marshall, Fink (2002). *Brain*, 125, 1024-1038.* | 1.5T fMRI | 11 | -- | Semantic fluency relative to overlearned sequence fluency (days of the week) | 1 | -- | YA: 32<br>OA: N/A |
| Noppeney & Price (2002). *NeuroImage*, 15, 927-935.* | PET | 12 | -- | Semantic decision task | 2 | -- | YA: 24 (20-30)<br>OA: N/A |
| McDermott, Petersen, Watson, Ojemann (2003). *Neuropsychologia*, 41, 293-303.* | 1.5T fMRI | 20 | -- | Word lists of semantic compared to phonological | 3 | -- | YA: 22.1 (18-32)<br>OA: N/A |
| Xiang, Lin, Ma, Zhang, Bower, Weng, Gao (2003). *Hum Brain Map*, 18, 208-214.* | 1.5T fMRI | 6 | -- | Semantic discrimination | 1 | -- | YA: 21-36<br>OA: N/A |
| Tieleman, Seurinck, Deblaere, Vandemaële, Vingerhoets, Achten (2005). *NeuroImage*, 26, 565-572.* | 1.5T fMRI | 22 | -- | Semantic compared to perceptual categorization | 3 | -- | YA: 29 (22-47)<br>OA: N/A |
| Frings, Dimitrova, Schorn, Elles, Hein-Kropp, Gizewski, Diener, Timmann (2006). *Neurosci Letters*, 409, 19-23.* | 1.5T fMRI | 16 | -- | Verb generation task | 3 | -- | YA: 24.9 (18-35)<br>OA: N/A |
| Callan, Tsytsarev, Hanakawa, Callan, Katsuhara, Fukuyama, Turner (2006). *NeuroImage*, 31, 1327-1342.** | 3T fMRI | 16 | -- | Listening to and covert production of singing relative to speech | 2 | -- | YA: 26 (19-47)<br>OA: N/A |
| Sweet, Paskavitz, Haley, Gunstad, Mulligan, Nyalakanti, Cohen (2008). *Neuropsychologia*, 46, 1114-1123.** | 1.5T fMRI | 34 | -- | Phonological similarity during verbal working memory | 1 | -- | YA: 37.24 (18-80)<br>OA: N/A |
| Durisko & Fiez (2010). *Cortex*, 46, 896-906.** | 3T fMRI | 19 | -- | Delayed serial recall task with letters | 2 | -- | YA: 23 (19-33)<br>OA: N/A |
| Davis, Kragel, Madden, Cabeza (2012). *Cereb Cortex*, 22, 232-242. | 3T fMRI | 18 | 16 | Semantic matching task | 0 | 1 | YA: 21.70<br>OA: 68.06 |
| Shafit, Randall, Stamatakis, Wright, Tyler (2012). *J Cognitive Neurosci*, 24, 1434-1446. | 3T fMRI | 14 | 16 | Lexical decision task, imageability of high and low competition words | 2 | 0 | YA: 23.86<br>OA: 75.75 |

**Table 3.** Included studies of motor tasks. Notably, while many of the included studies had both YA and OA, in some instances OA data came from clinical studies wherein the OA served as a control group. Further, additional YA data came from studies included in prior meta-analyses. ^#^Studies included only as part of Stoodley & Schmahmann, 2009. *Studies included as part of Stoodley & Schmahmann, 2009 and E et al., 2014. **Studies only included in E et al., 2014. “—” denotes studies where group a particular age group was not included and as such no coordinates are possible. Cases where there were no cerebellar coordinates are indicated by a 0 in the appropriate foci column. Note: the study by Linortner and colleagues (2012) is listed twice. Because two distinct and unique samples of older adults were included, the foci were entered separately into the analyses. Huang and colleagues (2010) looked at older adults in a separate experiment investigating working memory (see Table 4 below), and the motor task was only conducted in the young adult sample. However, the data are included here as based on our search terms, this study met our criteria. N/A: not applicable.

| Study | Imaging Modality | N, YA | N, OA | Task | # YA Foci | # OA Foci | Age: Mean (Range) |
| --- | --- | --- | --- | --- | --- | --- | --- |
| **Motor** |  |  |  |  |  |  |  |
| Wurster, Graf, Ackermann, Groth, Kassubek, Riecker (2015). *Brain Struct Func*, 220, 1637-1648. | 3T fMRI | -- | 10 | Finger tapping | -- | 2 | YA: N/A<br>OA: 64.9 |
| Blumen, Holtzer, Brown, Gazes, Verghese (2015). *Hum Brain Mapp*, 35, 4090, 4104. | 3T fMRI | -- | 33 | Imagined walking and walking while talking | -- | 1 | YA: N/A<br>OA: 73.03 |
| Allali, van der Meulen, Beauchet, Rieger, Vuilleumier, Assal (2014). *J Gerontol A Biol Sci Med Sci*, 69, 1389-1398. | 3T fMRI | 14 | 14 | Motor imagery | 1 | 3 | YA: 27.0<br>OA: 66.0 |
| Wittenberg, Lovelace, Foster, Maldjian (2014). *Brain Imaging Behav*, 8, 335-345. | 1.5T fMRI | 12 | 12 | Ecologically valid motor self-care tasks (e.g., buttoning & zipping) and finger tapping | 11 | 6 | YA: 29.0<br>OA: 61.0 |
| Crémers, D'Ostilio, Stamatakis, Delvaux, Garraux (2012). *Movement Disorders*, 27, 1498-1505. | 3T fMRI | -- | 15 | Imagined gait relative to imagined standing | -- | 3 | YA: N/A<br>OA: 63.8 |
| Vidoni, Thomas, Honea, Koskutova, Burns (2012). *J Neurol Phys Ther*, 36, 8-16. | 3T fMRI | -- | 10 | Power grip | -- | 3 | YA: N/A<br>OA: 73.6 |
| Zwergal, Linn, Xiong, Brandt, Strupp, Jahn (2012). | 3T fMRI | 20 | 20 | Imagined movement, standing and walking | 0 | 5 | YA: 24-40<br>OA: 60-78 |
| *Neurobiol Aging*, 33, 1073-1084. |  |  |  |  |  |  |  |
| Askim, Indredavik, Haberg (2010). *Arch Phys Med Rehabil*, 91, 1529-1536. | 1.5T fMRI | -- | 15 | Finger tapping – paced and self-paced | -- | 3 | YA: N/A<br>OA: 65.9 (50-75) |
| Allen & Humphreys (2009). *Current Biol*, 19, 1044-1049. | 3T fMRI | -- | 7 | Somatomotor tactile stimulation | -- | 1 | YA: N/A<br>OA: >74 |
| Eckert, Peschel, Heinze, Rotte (2006). *J Neurol*, 253, 199-207. | 1.5T fMRI | -- | 9 | Opening and closing fist | -- | 1 | YA: N/A<br>OA: 60.6 |
| Heuninckx, Wenderoth, Debaere, Peeters, Swinnn (2005). *J Neurosci*, 25, 6787-6796. | 3T fMRI | 12 | 12 | Coordinated hand and foot movements | 8 | 31 | YA: 22.4 (20-25)<br>OA: 64.8 (62-71) |
| Rowe, Stephan, Friston, Frackowiak, Lees, Passingham (2002). *Brain*, 125, 267-289. | Not Specified | -- | 12 | Motor sequence learning | -- | 3 | YA: N/A<br>OA: 62.0 |
| Kalpouzos, Garzón, Simikov, Heiland, Salami, Persson, Bäckman (2017). *Cereb Cortex*, 27, 3427-3436. | 3T fMRI | 22 | 15 | Motor imagery | 1 | 2 | YA: 36.8<br>OA: 69.7 |
| King, Saucier, Albouy, Fogel, Rumpf, Klann, Buccino, Binkofski, Classen, Karni, Doyon (2017). *Cereb Cortex*, 27, 1588-1601. | 3T fMRI | -- | 26 | Motor sequence learning, initial training only included here | -- | 3 | YA: N/A<br>OA: 63.5 |
| Wang, Qiu, Liu, Yan, Yang, Zhang, Zhang, Sang, Zheng (2014). *Neuroreport*, 56, 339-348. | 3T fMRI | 19 | 20 | Motor execution and imagery | 6 | 8 | YA: 36.5 (20-23)<br>OA: 62.5 (52-82) |
| Heuninckx, Wenderoth, Swinnen (2010). *Neurobiol Aging*, 31, 301-314. | 3T fMRI | 12 | 12 | Externally and internally guided movements | 6 | 8 | YA: 23.5 (21-27)<br>OA: 66.9 (63-73) |
| Riecker, Gröschel, Ackermann, Steinbrink, Witte, Kastrup (2006). *NeuroImage*, 32, 1345- | 1.5T fMRI | 10 | 10 | Motor tapping | 1 | 1 | YA: 23.0 (18-26)<br>OA: 66.0 (58-82) |
| 1354. |  |  |  |  |  |  |  |
| Zapparoli, Invernizzi, Gandola, Verardi, Berlingeri, Sberna, De Santis, Zerbi, Banfi, Bottini, Paulesu (2012). *Exp Brain Res*, 224, 519-540. | 1.5T fMRI | 24 | 24 | Finger to thumb opposition | 1 | 0 | YA: 27.0<br>OA: 60.0 |
| Jäncke, Loose, Lutz, Specht, Shah (2000). *Cog Brain Res*, 10, 51-66.# | 1.5T fMRI | 8 | -- | Finger tapping | 3 | -- | YA: 20-32<br>OA: N/A |
| Lutz, Specht, Shah, Jäncke (2000). *NeuroReport*, 11, 1301-1306.# | 1.5T fMRI | 10 | -- | Finger tapping | 4 | -- | YA: 24.1 (21-29)<br>OA: N/A |
| Riecker, Wildgruber, Mathiak, Grodd, Ackermann (2003). *NeuroImage*, 18, 2003, 731-739.# | 1.5T fMRI | 8 | -- | Finger tapping | 12 | -- | YA: 23.75 (19-32)<br>OA: N/A |
| Hanakwa, Dimyan, Hallett (2008). *Cereb Cortex*, 18, 2775-2788.# | 3T fMRI | 13 | -- | Finger Tapping | 2 | -- | YA: 30.0 (21-48)<br>OA: N/A |
| Brunne, Skouen, Ersland, Grüner (2014). *Neurorehab Neral Repair*, 28, 874-884. | 3T fMRI | -- | 18 | Observation and execution of bimanual movements | -- | 4 | YA: N/A<br>OA: 60.6 |
| Taniwaki, Okayama, Yoshiura, Togao, Nakamura, Yamsaki, Ogata, Shigeto, Shyagi, Kira, & Tobimatsu (2007). *NeuroImage*, 36, 1263-1276. | 1.5T fMRI | 12 | 12 | Externally triggered or self-initiated finger movements | 6 | 4 | YA: 24.9 (23-29)<br>OA: 62.9 (53-72) |
| Taniwaki, Yoshiura, Ogata, Togao, Yamashita, Kida, Miura, Kira, Tobimatsu (2013). *Brain Res*, 1512, 45-59. | 1.5T fMRI | -- | 12 | Externally triggered or self-initiated finger movements | -- | 4 | YA: N/A<br>OA: 62.0 (54-72) |
| Daselaar, Rombouts, Veltman, Raaijmakers, Jonker (2003). *Neurobiol* | 1.5T fMRI | 26 | 40 | Motor sequence learning | 1 | 3 | YA: 32.4 (30-35)<br>OA: 66.4 (63-71) |
| *Aging*, 24, 1013-1019. |  |  |  |  |  |  |  |
| Onozuka, Fujita, Watanabe, Hirano, Niwa, Nishiyama, Saito (2003). *J Dent Res*, 82, 657-660. | 1.5T fMRI | 10 | 10 | Chewing | 1 | 1 | YA: 19-26<br>OA: 65-73 |
| Rijntjes, Buechel, Kiebel, Weiller (1999). *NeuroReport*, 10, 3653-3658.# | 2T fMRI | 9 | -- | Finger flexion and extensions | 3 | -- | YA: 32.0<br>OA: N/A |
| Hankawa, Immisch, Toma, Dimyan, van Gelderen, Hallett (2003). *J Neurophysiol*, 89, 989-1002.# | 1.5T fMRI | 10 | -- | Finger tapping | 3 | -- | YA: 32.0<br>OA: N/A |
| Loibl, Beutling, Kaza, Lotze (2011). *Behav Brain Res*, 223, 280-286. | 1.5T fMRI | 18 | 17 | Passive wrist movement, fist clenching, precision grip | 21 | 17 | YA: 25.39 (23-30)<br>OA: 66.65 (57-72) |
| Linortner, Fazekas, Schmidt, Ropele, Pendl, Petrovic, Loitfelder, Neuper, Enzinger (2012). *Neurobiol Aging*, 197, e1-191-e9-17. | 3T fMRI | -- | 17 | Ankle and finger movements | -- | 2 | YA: N/A<br>OA: 63.59 (48-84) |
| Linortner, Fazekas, Schmidt, Ropele, Pendl, Petrovic, Loitfelder, Neuper, Enzinger (2012). *Neurobiol Aging*, 197, e1-191-e9-17. | 3T fMRI | -- | 13 | Ankle and finger movements | -- | 3 | YA: N/A<br>OA: 73.31 (48-84) |
| Kim, Lee, Lee, Song, Yoo, Lee, Kim, Chang (2010). *Neurological Res*, 32, 995-1001. | 3T fMRI | 20 | 26 | Weighted and unweighted elbow flexion/extension | 0 | 3 | YA: 23.0<br>OA: 65.5 |
| Rieckman, Fischer, Bäckman (2010). *NeuroImage*, 50, 1303-1312. | 1.5T fMRI | 14 | 13 | Serial reaction time task | 2 | 1 | YA: 24.71<br>OA: 68.08 |
| Huang, Lee, Hsiao, Kuan, Wai, Ko, Wan, Hsu, Liu (2010). *J Neurosci Methods*, 189, 257-266. | 1.5T fMRI | 16 | -- | Hand flexion | 1 | -- | YA: 22.0 (18-25)<br>OA: N/A |
| Bo, Peltier, Noll, Seidler (2011). *Neurosci Letters*, 504, | 3T fMRI | 14 | 14 | Motor sequence learning with symbolic and spatial | 1 | 0 | YA: 21.4<br>OA: 72.7 |
| 68-72. |  |  |  | presentation of stimuli |  |  |  |
| Dennis, Cabeza (2011). *Neurobiol Aging*, 2318.17-2318.e30. | 4T fMRI | 12 | 12 | Serial reaction time task | 1 | 0 | YA: 22.2 (18-30)<br>OA: 67.4 (60-79) |

**Table 4.**
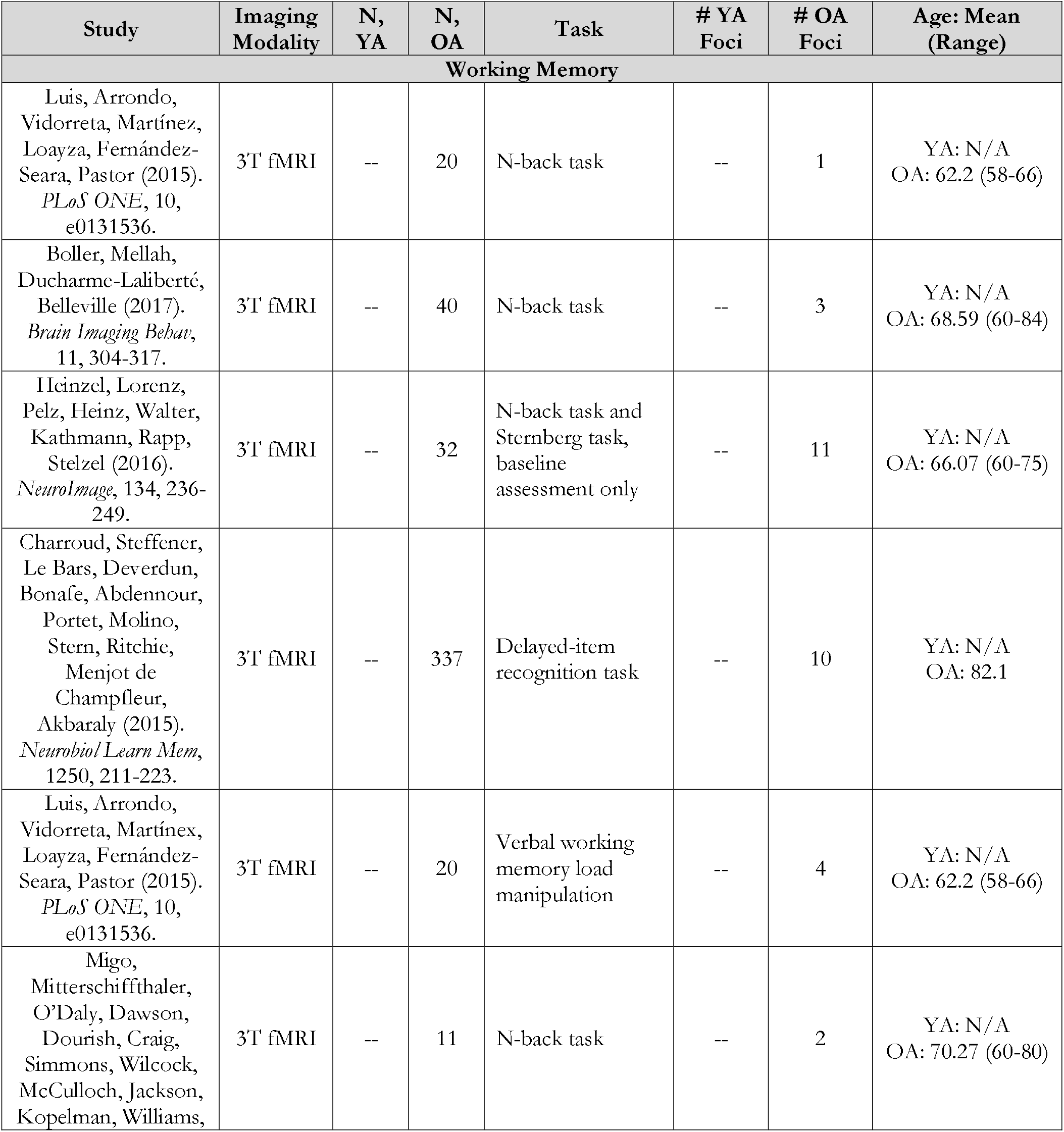
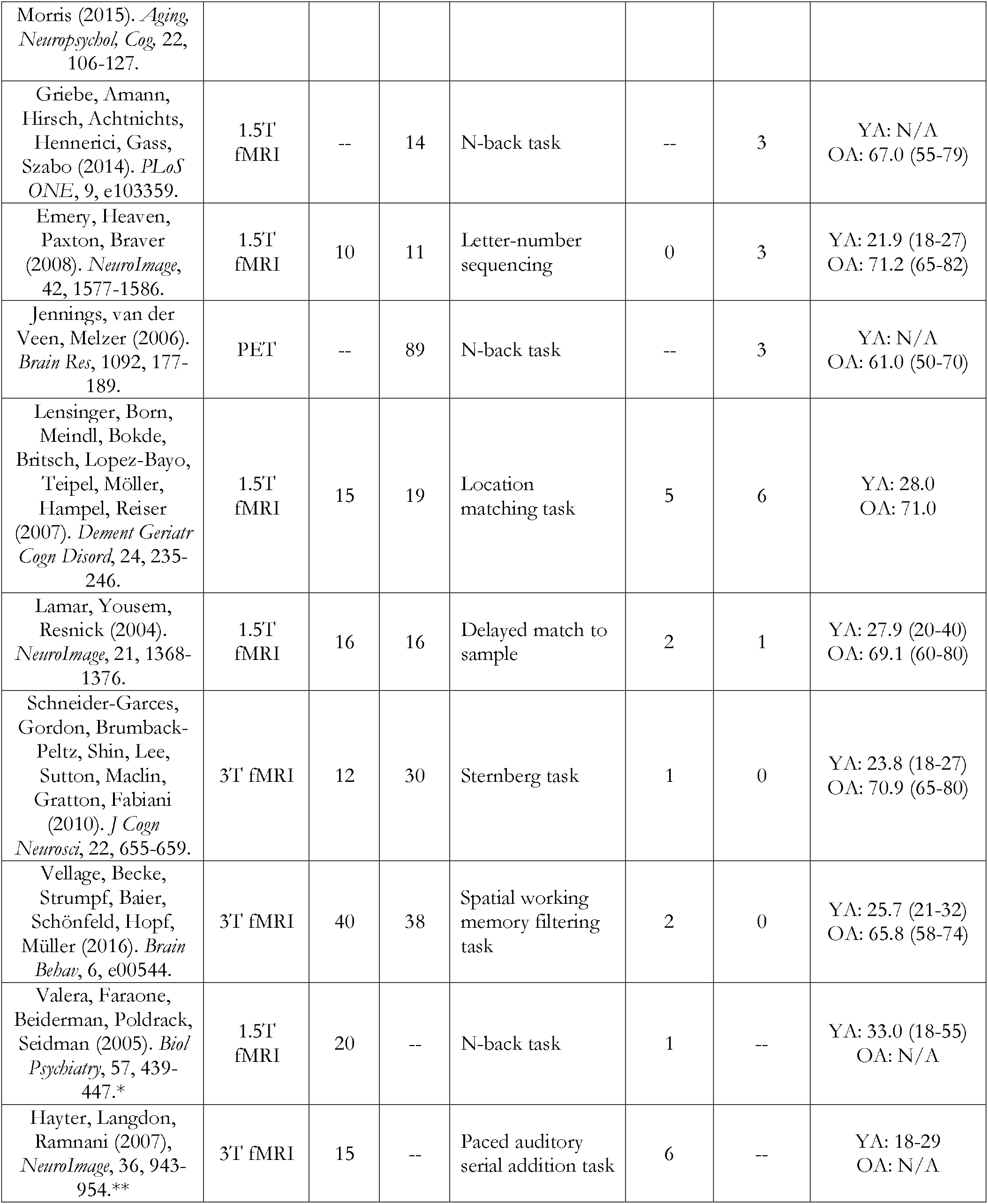
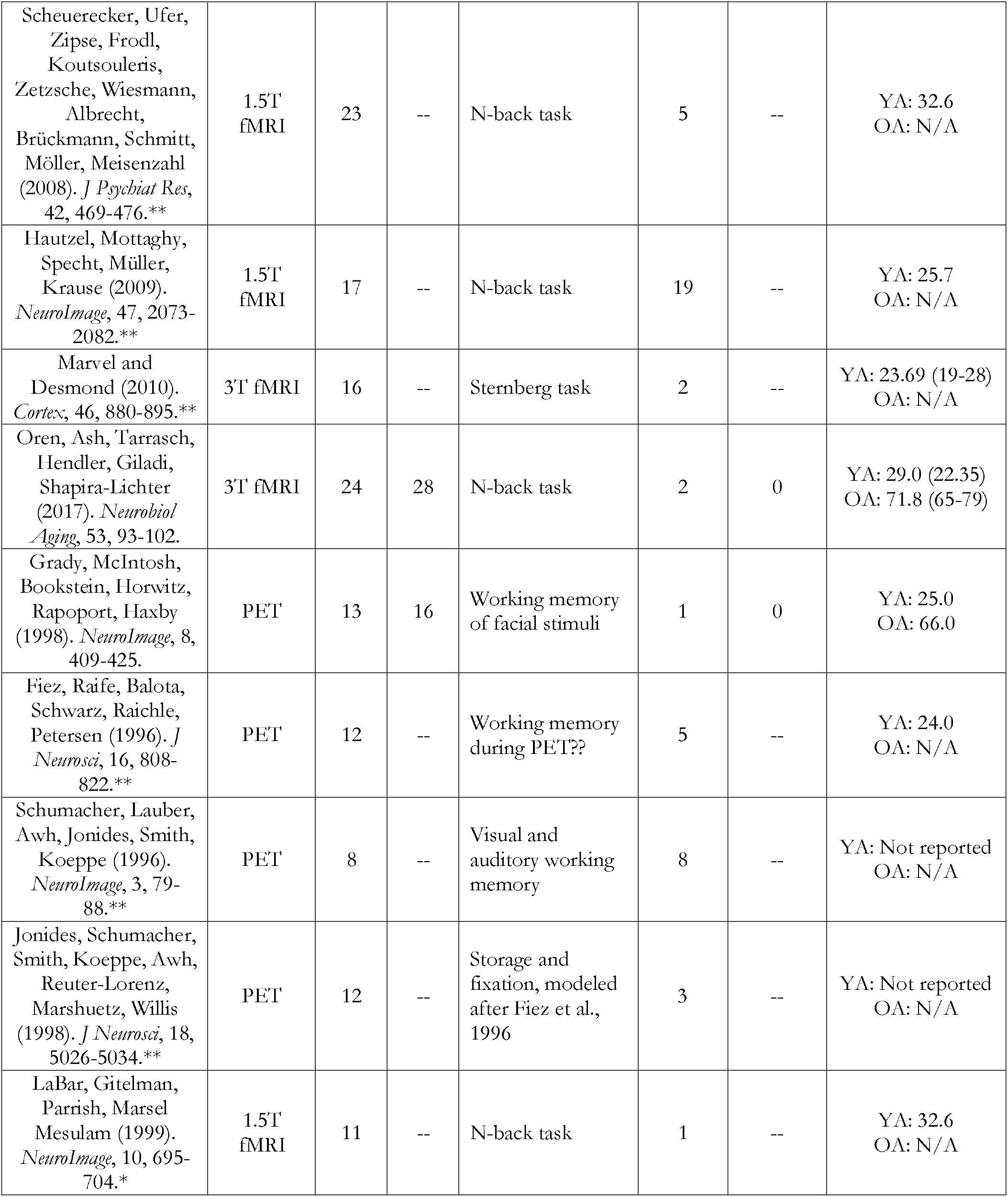
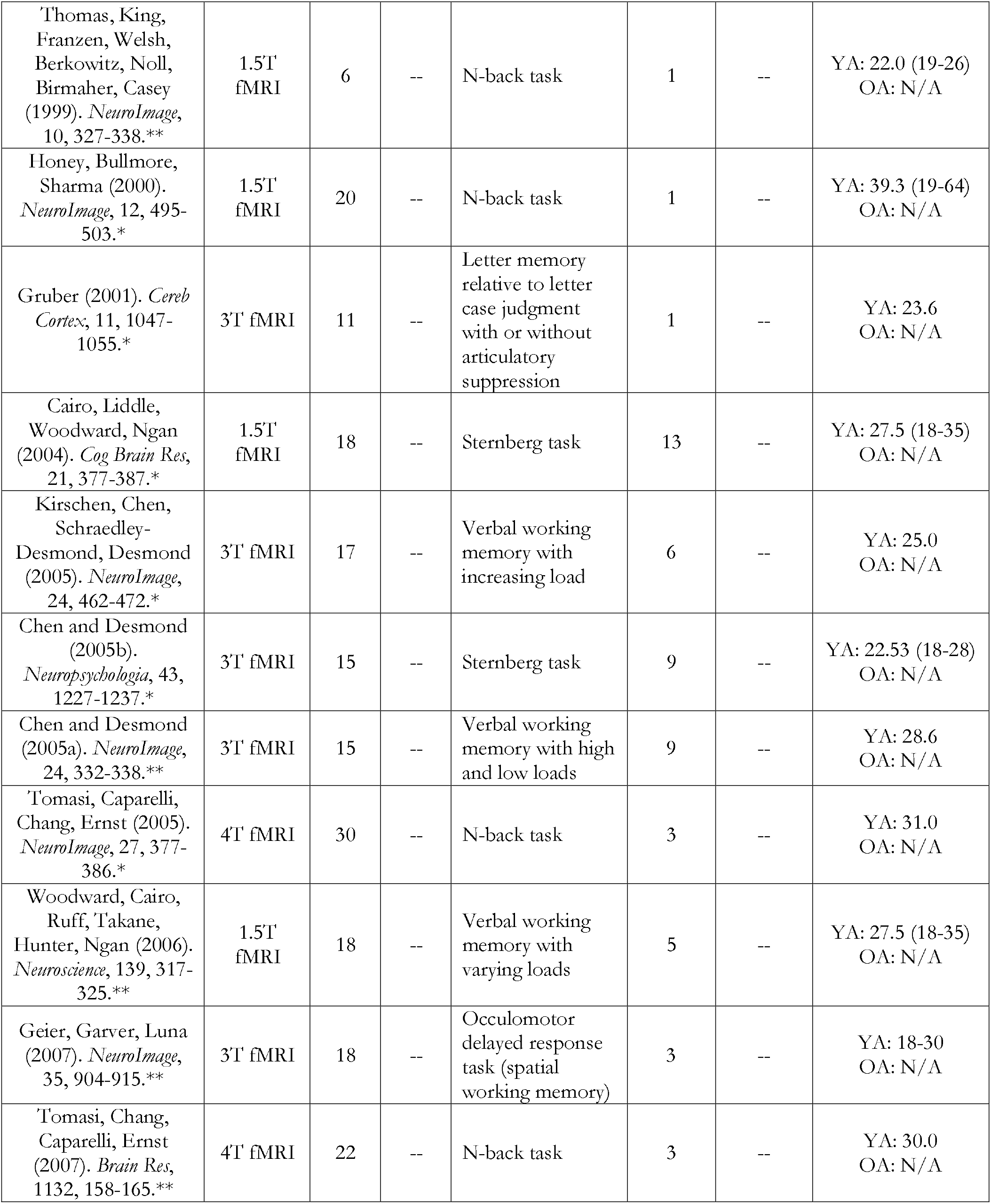
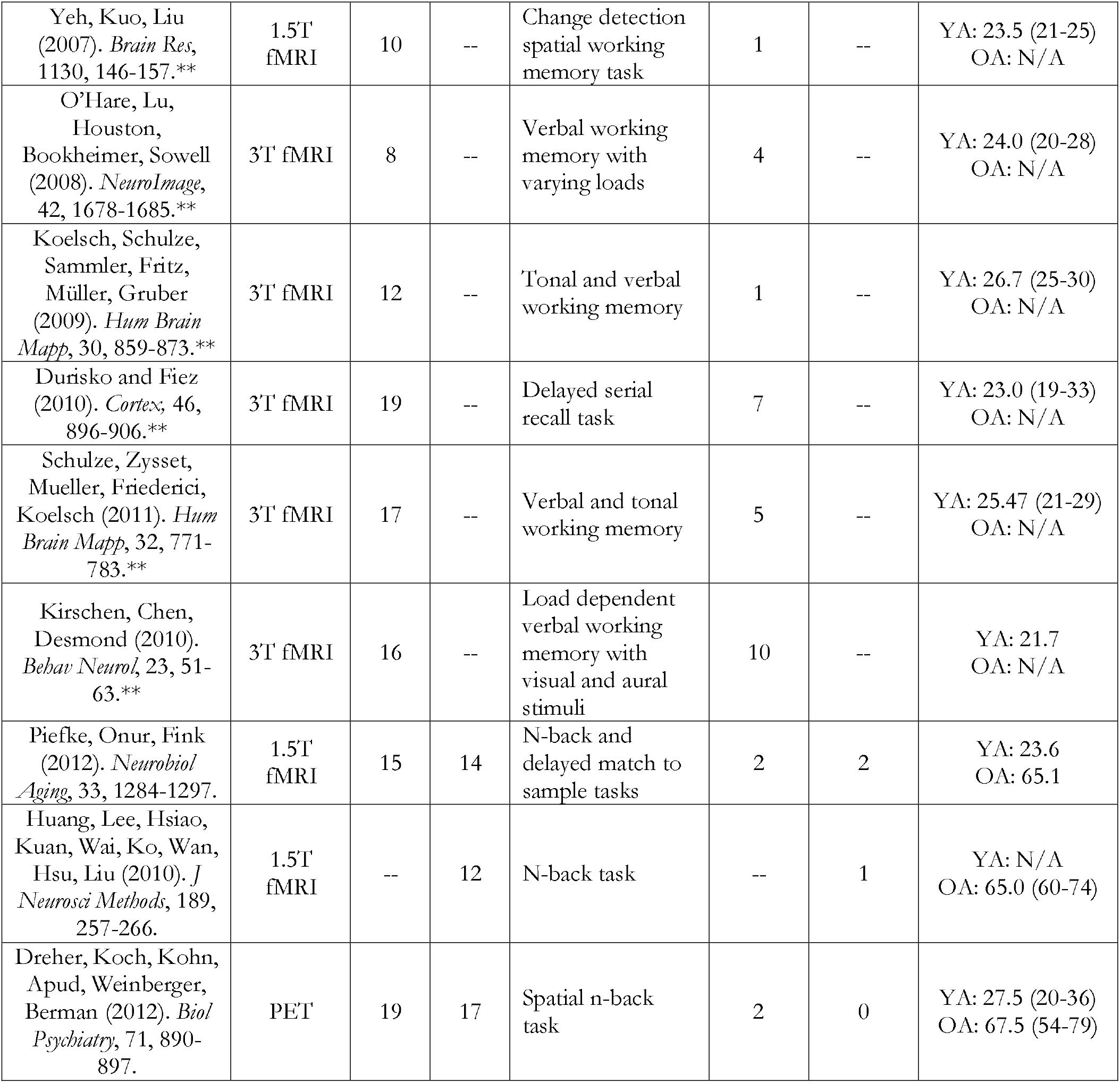
Included studies of working memory. Notably, while many of the included studies had both YA and OA, in some instances OA data came from clinical studies wherein the OA served as a control group. Further, additional YA data came from studies included in prior meta-analyses. ^#^Studies included only as part of Stoodley & Schmahmann, 2009. *Studies included as part of Stoodley & Schmahmann, 2009 and E et al., 2014. **Studies only included in E et al., 2014. “—” denotes studies where group a particular age group was not included and as such no coordinates are possible. Cases where there were no cerebellar coordinates are indicated by a 0 in the appropriate foci column. N/A: not applicable.

**Table 5.** Included studies of long-term memory. Notably, while many of the included studies had both YA and OA, in some instances OA data came from clinical studies wherein the OA served as a control group. “—” denotes studies where group a particular age group was not included and as such no coordinates are possible. Cases where there were no cerebellar coordinates are indicated by a 0 in the appropriate foci column. N/A: not applicable

| Study | Imaging Modality | N, YA | N, OA | Task | # YA Foci | # OA Foci | Age: Mean (Range) |
| --- | --- | --- | --- | --- | --- | --- | --- |
| **Long-Term Memory** |  |  |  |  |  |  |  |
| Vidal-Piñeir, Martín-Trias, Arenaza-Urquijo, Sala-Llonch, Clemente, Mena-Sánchez, Bargalló, Falcón, Pascual-Leone, Bartrés-Faz (2014). *Brain Stim*, 7, 287-296. | 3T fMRI | -- | 24 | Memory for deep and shallow encoding | -- | 3 | YA: N/A<br>OA: 71.75 (61-80) |
| Beason-Held, Golski, Kraut, Esposito, Resnick (2005). *Neurobiol Aging*, 26, 237-250. | PET | -- | 11 | Verbal and figural encoding and recognition | -- | 10 | YA: N/A<br>OA: 71.1 (63-82) |
| Peira, Ziaei, Persson (2016). *NeuroImage*, 125, 745-755. | 3T fMRI | 15 | 15 | Prospective memory | 0 | 2 | YA: 22.4 (20-26)<br>OA: 68.1 (64-74) |
| Cohen, Rissman, Suthana, Castel, Knowlton (2016). *NeuroImage*, 125, 1046-1062. | 3T fMRI | -- | 23 | Memory for words with high and low value | -- | 3 | YA: N/A<br>OA: 68.7 (60-80) |
| Gong, Fu, Wang, Franz, Long (2014). *Int'l J Aging Hum Dev*, 79, 23-54. | 3T fMRI | -- | 12 | Emotional autobiographical memory retrieval | -- | 8 | YA: N/A<br>OA: 66.3 (60-74) |
| Miotto, Balardin, Savege, Martin, Batistuzzo, Amaro, Nitrini (2014). *Arq Neuropsiquiatr*, 72, 663-670. | 3T fMRI | -- | 17 | Memory for semantically related and unrelated words | -- | 1 | YA: N/A<br>OA: 68.12 |
| Brassen, Büchel, Weber-Fahr, Leimbek, Sommer, Braus (2009). *Neurobiol Aging*, 30, 1147-1156. | 3T fMRI | 14 | 14 | Correct retrieval during verbal episodic memory | 0 | 1 | YA: 25.6 (21-33)<br>OA: 64.9 (60-71) |
| Bartrés-Faz, Serra-Grabulosa, Sun, Solé-Padullés, Rami, Molineuvo, Bosch, Mercader, Bargalló, Falcón, vendrell, Junqué, D'Esposito (2008). *Neurobiol Aging*, 29, 1644-1653. | 1.5T fMRI | -- | 20 | Encoding of face-name pairs | -- | 1 | YA: N/A<br>OA: 66.0 |
| Maguire and Frith (2003). *Brain*, 126, 1511-1523. | 2T fMRI | 12 | 12 | Autobiographical memory | 4 | 4 | YA: 32.42 (23-39)<br>OA: |
| Gronholm, Rinne, Vorobyev, Laine (2005). *Cog Brain Res*, 25, 359-371. | PET | -- | 10 | Memory for learned objects | -- | 6 | YA: N/A<br>OA: 74.75 (67.80) |
| Milton, Butler, Benattayallah, Zeman (2012). *Neuropsychologia*, 50, 3528-3541. | 1.5T fMRI | -- | 17 | Long-term autobiographical memory | -- | 6 | YA: N/A<br>OA: >50 |
| Lam, Wächter, Globas, Karnath, Luft (2013). *Hum Brain Mapp*, 34, 176-185. | 3T fMRI | -- | 10 | Weather prediction task | -- | 2 | YA: N/A<br>OA: 64.6 (43-85) |
| Antonova, Parslow, Brammer, Dawson, Jackson, Morris (2009). *Memory*, 17, 125-143. | 1.5T fMRI | 10 | 10 | Long-term memory of spatial information | 4 | 4 | YA: 23.6 (20-26)<br>OA: 72.14 (64-79) |
| Dannhauser, Shergill, Stevens, Lee, Seal, Walker, Walker (2008). *Cortex*, 44, 869-880. | 1.5T fMRI | -- | 10 | Verbal episodic memory | -- | 1 | YA: N/A<br>OA: 68 (50-84) |
| Bowman and Dennis (2015). *Brain Res*, 1612, 2-15. | 3T fMRI | 17 | 22 | Remember/know judgments of long-term memory | 0 | 1 | YA: 21.28 (18-25)<br>OA: 74.18 (67-83) |
| Kircher, Weis, Leube, Freymann, Erb, Jessen, Grodd, Heun, Krach (2008). *Eur Arch Psychiatry Clin Neurosci*, 258, 363-372. | 1.5T fMRI | -- | 29 | Subsequent memory effect | -- | 1 | YA: N/A<br>OA: 67.7 (60-81) |
| Daselaar, Veltman, Rombouts, Raaijmakers, Jonker (2003). *Brain*, 126, 43-56. | 1.5T fMRI | 17 | 19 | Correct rejection or recognition at retrieval | 3 | 2 | YA: 32.7 (30-35)<br>OA: 66.4 (63-71) |
| Haist, Gore, Mao (2001). *Nature Neurosci*, 4, 1139-1145. | 1.5T fMRI | -- | 8 | Remote memory for famous faces | -- | 2 | YA: N/A<br>OA: 64.6 (60-70) |
| Iidaka, Sadato, Yamada, Murata, Omori, Yonekura (2001). *Cog Brain Res*, 11, 1-11. | 1.5T fMRI | 7 | 7 | Pictorial information, abstract object encoding | 0 | 1 | YA: 25.7<br>OA: 66.2 |
| Madden, Turkington, Provenzale, Denny, Hawk, Gottlob, Coleman (1999). *Hum Brain Map*, 7, 115-135. | PET | 12 | 12 | Recognition memory task | 1 | 2 | YA: 23.17 (20-29)<br>OA: 71.0 (62-79) |
| Gao, Cheung, Chan, Chu, Mak, Lee (2014). *PLoS ONE*, 9, e90307. | 3T fMRI | 13 | 13 | Prospective memory | 1 | 0 | YA: 27.1<br>OA: 76.2 |
| Grady, McIntosh, Horwitz, Maisog, Ungerleider, Mentis, Pietrini, Schapiro, Haxby (1995). *Science*, 269, 218-221. | PET | 10 | 10 | Recognition compared to matching in long-term memory | 3 | 0 | YA: 25.2<br>OA: 69.4 |
| Zamboni, de Jager, Drazich, Douaud, Jenkinson, Smith, Tracey, Wilcock (2013). *Neurobiol Aging*, 34, 961-972. | 3T fMRI | -- | 28 | Paired associates task | -- | 1 | YA: N/A<br>OA: 74.4 (64-91) |
| Braskie, Small, Bookheimer (2009). *Hum Brain Map*, 30, 3981-3992. | 3T fMRI | -- | 32 | Long term memory of word lists at retrieval | -- | 2 | YA: N/A<br>OA: 60.0 (42-77) |
| Düzel, Scütze, Yonelinas, Heinze (2011). *Hippocampus*, 21, 803-814. | 1.5T fMRI | 24 | 56 | Incidental encoding task, activation at recollection | 4 | 0 | YA: 23.0<br>OA: 65.0 |

To complete our analyses of age differences, we used the YA control samples from the OA literature, as opposed to doing an additional search focused on YA alone. To date, there have been several meta-analyses investigating task activation patterns in both the motor and cognitive domains in healthy YA (Bernard & Seidler, 2013a; E, Chen, Ho, & Desmond, 2012; Stoodley & Schmahmann, 2009), and such an analysis would be redundant, and is beyond the scope of the present investigation. Furthermore, we were concerned about considerable differences in the sample sizes that would potentially bias our group comparison analyses, as the YA literature is substantially larger. However, because many of the studies in our OA sample included OA that served as controls for an age-related disease group, we had a limited sample of YA studies. To better equate our groups, we included all the studies and foci from prior meta-analyses investigating cerebellar functional activation (E et al., 2012; Stoodley & Schmahmann, 2009) in overlapping task categories. Additional motor foci were extracted from the studies included by Stoodley & Schmahmann (2009), while those for language, working memory, and cognitive function (categorized as executive function) were taken from E and colleagues (2012). Notably, E and colleagues (2012) also had substantial overlap with Stoodley & Schmahmann (2009) as they included all of the papers from the prior analysis, as well as new additions to the literature. Long-term memory was not included in these prior investigations. Notably, we did not gather additional YA papers to match the OA papers for two reasons. First, though we could match based on task type, we were concerned that this could introduce selection bias. For many tasks there are multiple papers that have similar sample sizes as existing OA studies and additional inclusion criteria could have been biased. Second, though additional YA papers may have cerebellar foci, there would be no guarantee that the number of foci would be matched across studies even when taking this approach.

The literature searches and initial inclusion decisions were completed by T.M., A.D.N., Y.L., J.R.M.G., H.K.B., and H.K.H. Inclusion was confirmed and coordinates for each study were checked, prior to analysis, by J.A.B. After scanning the literature and the inclusion of the studies from both Stoodley & Schamhmann (2009) and E and colleagues (2012), we had 175 studies, including data from 1,710 YA (403 foci) and 2,160 OA (307 foci) individuals, concatenated across all task domains. Figure 1 provides a flow-chart demonstrating our paper screening procedure and exclusion regions, broadly defined. The initial search, and secondary broader aging search are presented separately.

### Activation Likelihood Estimation (ALE) Meta-Analysis

All analyses were completed using BrainMap GingerALE 3.0.2 (http://brainmap.org; Eickhoff, Bzdok, Laird, Kurth, & Fox, 2012; Eickhoff et al., 2009; Turkeltaub et al., 2012). ALE allows us to combine across studies, sites, scanning modalities, and study designs to investigate overlap in activation patterns, and the algorithm includes a metrics to account for variability in subjects and testing sites (Eickhoff et al., 2012). Unlike behavioral meta-analyses, because the algorithm looks at activation foci, and models these to account for uncertainty, the variability in design and analysis approaches can be reasonably accounted for. Foci were first organized for analysis by task domain; however, we also performed additional analyses concatenating across all cognitive task domains. As there are two standard atlas spaces used for normalization and presentation of activation (Montreal Neurological Institute (MNI) or Talairach) it is critical to ensure that all foci are in the same atlas space prior to analysis for the purpose of comparison across studies. As such, all foci in Talairach space were converted to MNI space prior to analysis. For studies where data were normalized directly to Talairach space, and those that specified the use of the Lancaster transform (icbm2tal; Lancaster et al., 2007), we used this transform to move them to MNI space. This transform was also used for studies published after the icbm2tal transform became available, and for which no specific transform information was provided. For studies where the Brett transform (mni2tal) was used to bring data from MNI space to Talairach space, and for articles published prior to 2007 without any transform details, we used the inverse Brett transform to bring the data into MNI space. All transforms were completed using tools available in GingerALE.

Once in MNI space, all activation foci were organized into text files for analysis with GingerALE. The ALE algorithm computes ALE values for all of the voxels in the brain, producing an estimation of the likelihood that a particular voxel is active under particular task conditions (Eickhoff et al., 2009). During analysis, GingerALE uses a full-width half maximum (FWHM) gaussian blur on each set of foci, with the size based off of the sample size used to generate each set of foci (Eickhoff et al., 2009). Output of our analyses indicated that the FWHM blur ranged from 8.46 to 11.37mm, across all analyses. In completing our analyses, we used the smaller more conservative mask option available in GingerALE, in conjunction with the non-additive ALE method (Turkeltaub et al., 2012). For within group analyses, all ALE maps were thresholded using a cluster-level family-wise error (FWE) p<.001 with 5,000 threshold permutations and a p-value of p<.001. Group contrasts and conjunctions were evaluated using an uncorrected p<.05 with 10,000 p-value permutations, and a minimum cluster size of 50 mm^3^. This approach is consistent with our prior meta-analyses (Bernard & Mittal, 2015; Bernard et al., 2017), as well as other recent work (e.g., Stawarczyk & D’Argembeau, 2015), and allows us to look at contrasts, even though GingerALE is not very robust when small numbers of studies (fewer than 15 per group) are used for group contrasts. The resulting areas of convergence from all analyses were localized using the Spatially Unbiased Infratentorial Template (SUIT) atlas (Diedrichsen, Balsters, Flavell, Cussans, & Ramnani, 2009). Foci located in the white matter in the area of the cerebellar nuclei were localized using an atlas of cerebellar nuclei (Dimitrova et al., 2002).

## Results

### Within Group Activation Convergence Across Studies

Because several meta-analyses have already been conducted investigating cerebellar activation across task domains in YA (Bernard & Seidler, 2013a; E et al., 2012; Stoodley & Schmahmann, 2009), we provide only a brief overview of the YA results. Details of the areas of activation overlap across studies for both age groups and each task domain are provided in Table 6 and presented visually in Figure 2. In YA, the motor and working memory analyses replicated prior meta-analyses investigating patterns of cerebellar functional activation (Stoodley & Schmahmann, 2009; E et al., 2012; Bernard & Seidler, 2013a), though notably, there was substantial overlap in the foci used for the analyses. Activation overlap across language tasks was also consistent with prior work with a large cluster extending across Crus I and Lobule VI, while that for other cognitive tasks, which primarily included executive function tasks, also paralleled prior work (Stoodley & Schmahmann, 2009; E et al., 2012). Notably, this area is also consistent with recent work mapping function in the cerebellum by King and colleagues (2019), where tasks similar to those categorized here showed activation in lateral posterior cerebellum. Finally, we extended prior meta-analyses with the inclusion of long-term memory. In YA, our results demonstrate activation overlap across tasks in Lobule VI and Crus I. The overlap in Crus I is consistent with Crus I activation seen with autobiographical recall by King and colleagues (2019).

**Figure 2.**
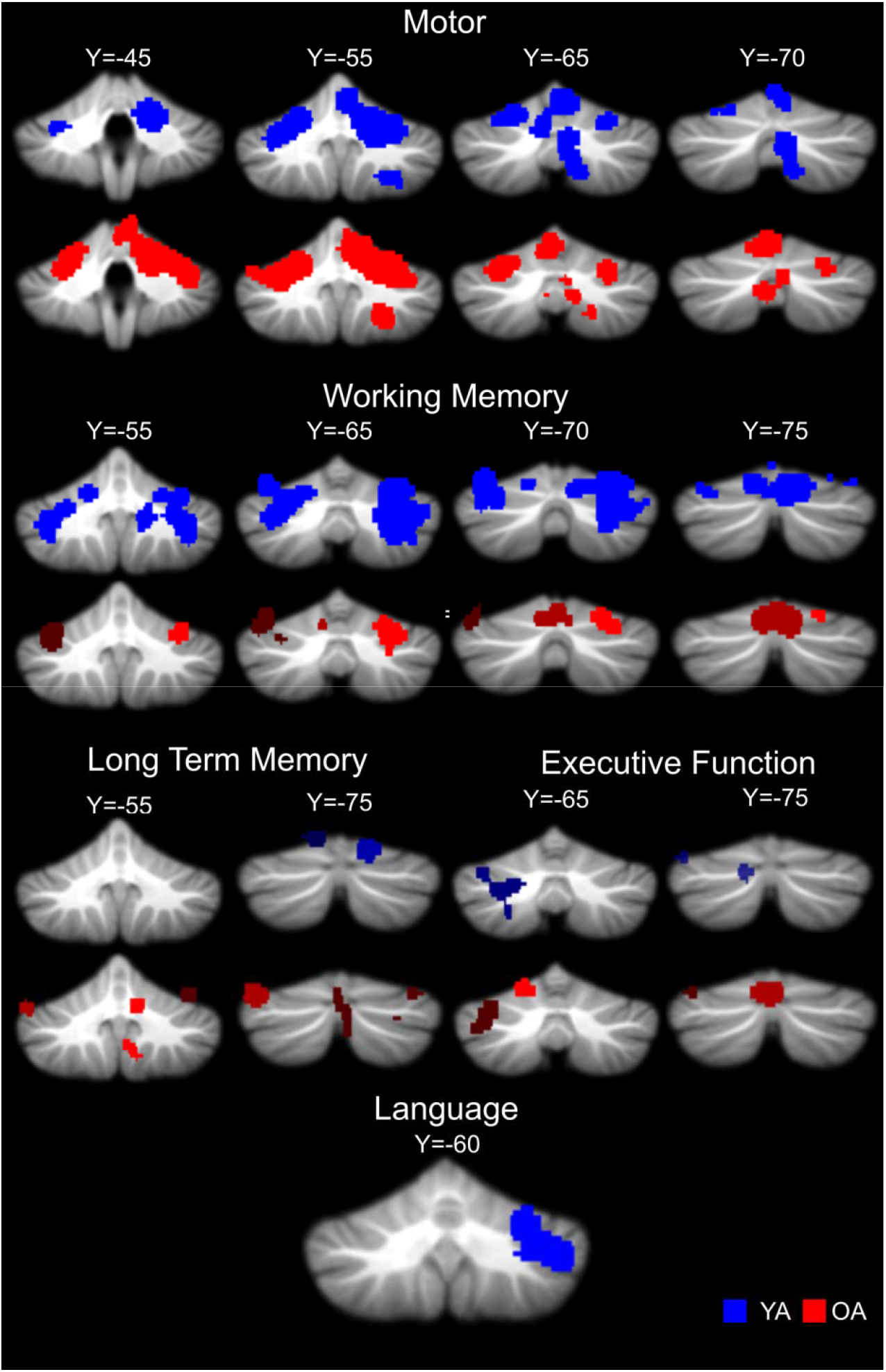
Activation overlap in the cerebellum across studies for each task domain in YA (blue) and OA (red). Areas of overlap are overlaid onto the SUIT cerebellum template. Notably, there were no significant areas of overlap across studies in OA for language tasks.

**Table 6.**
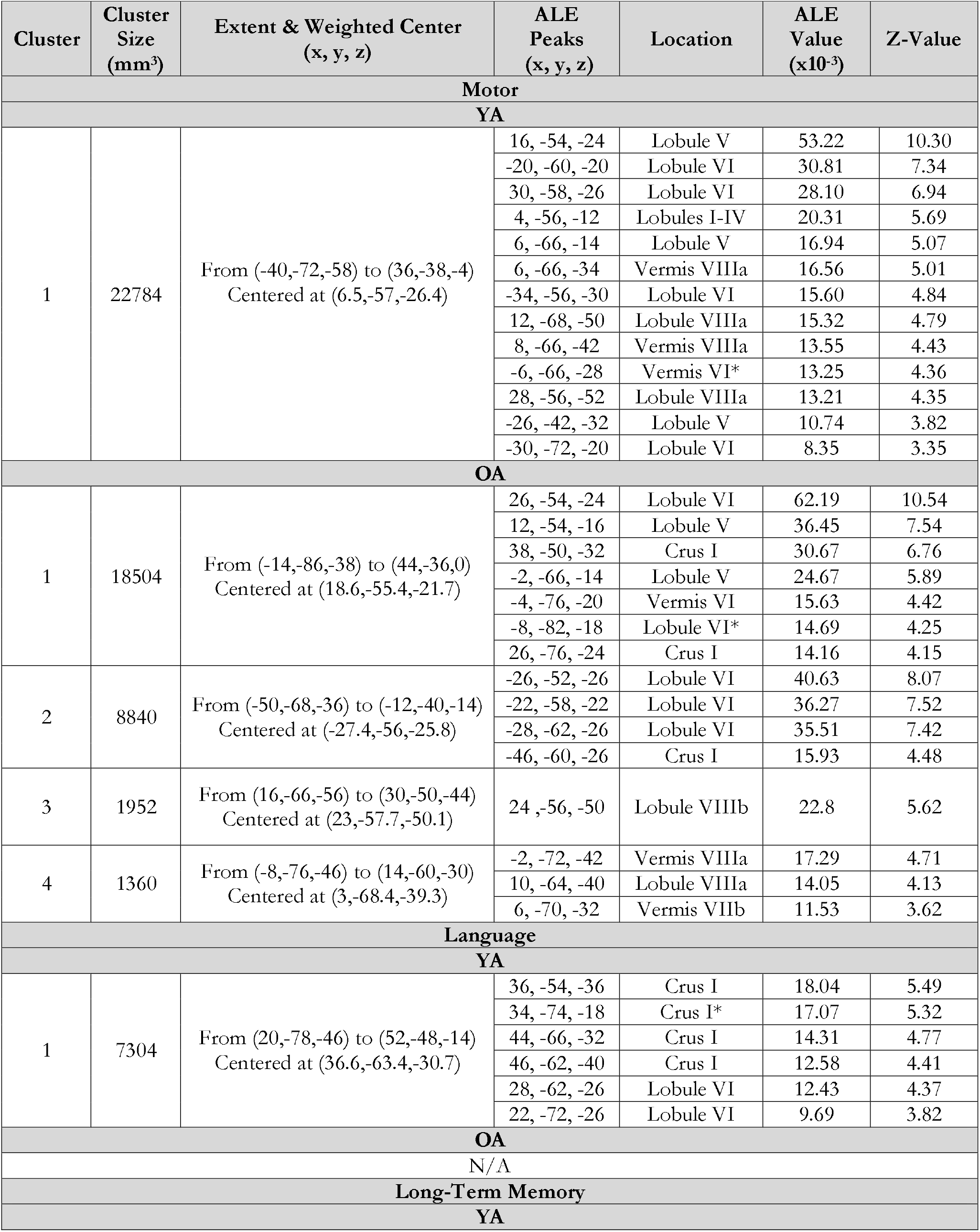
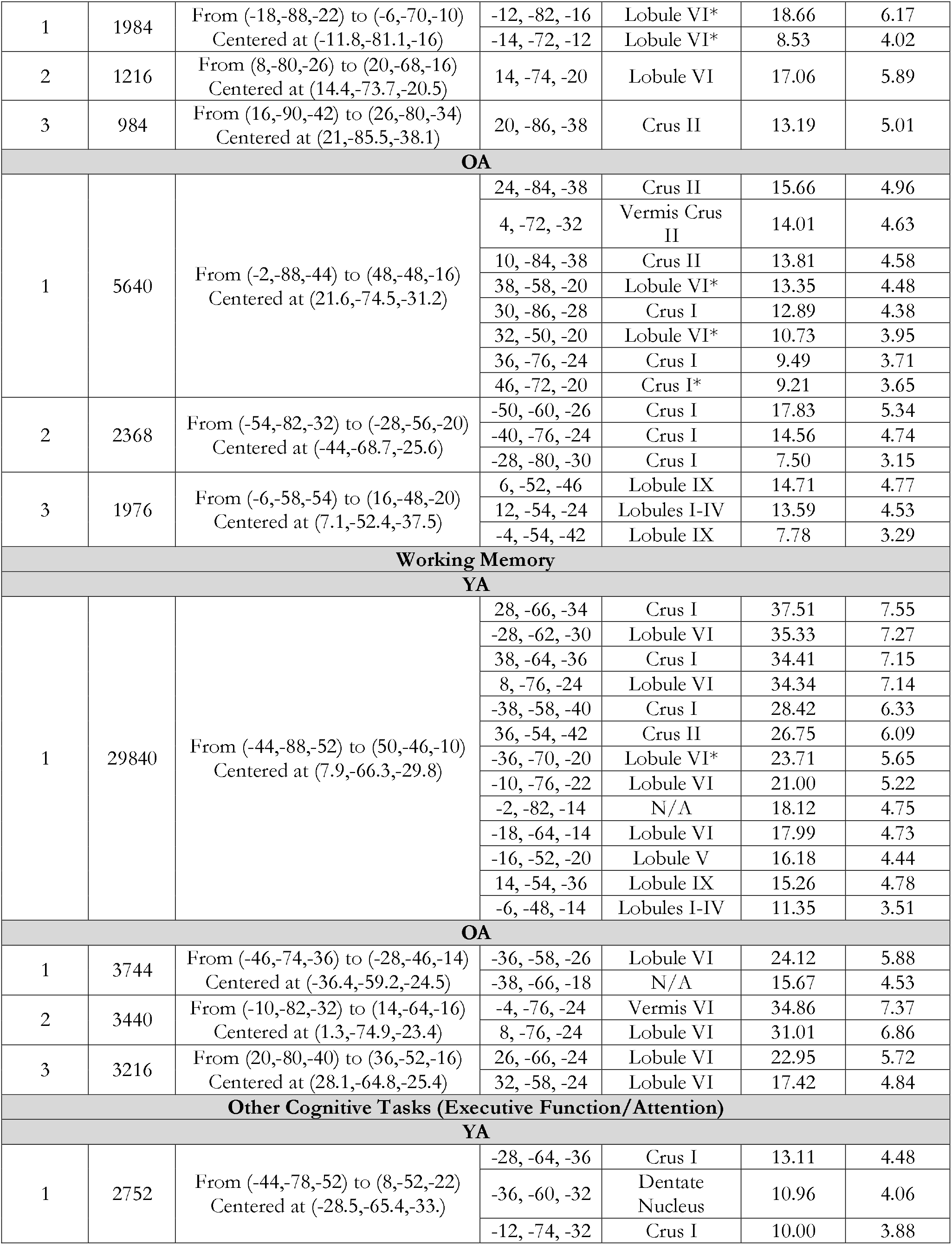
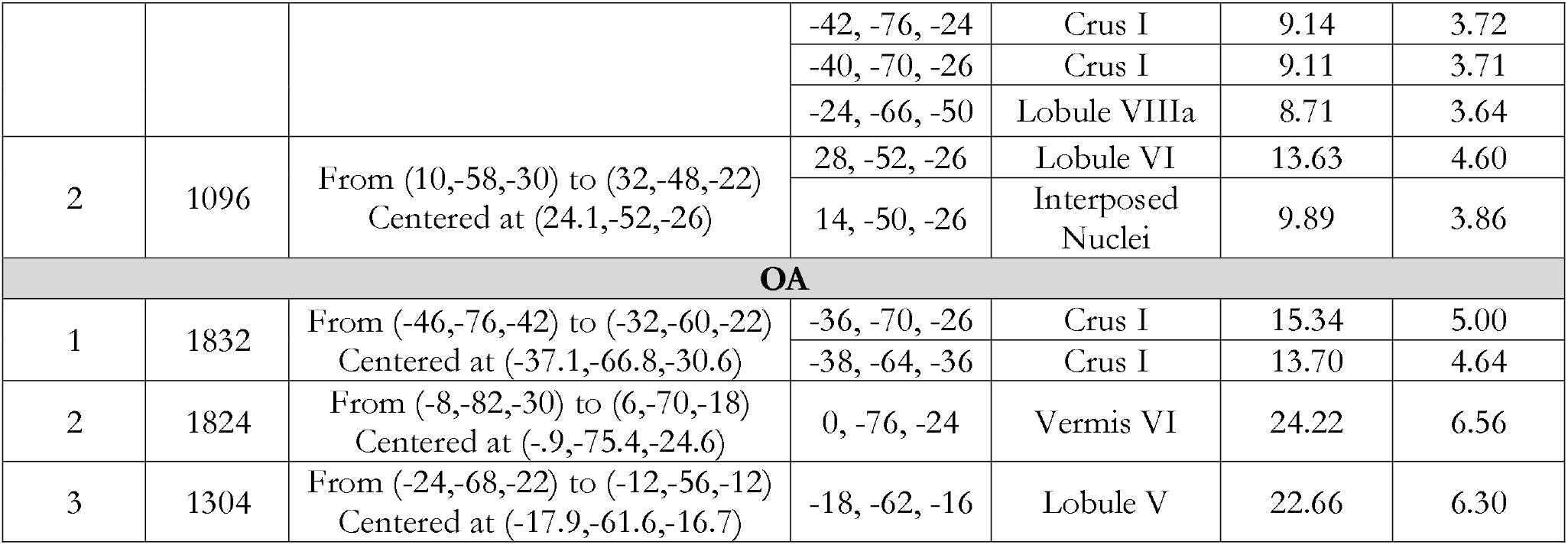
Activation by group and task. *Peak outside of SUIT Atlas space, the closest region to the reported peak is listed.

In OA, across studies motor task activation largely paralleled YA in the regions where we observed activation overlap. That is, activation was localized largely to the anterior cerebellum in lobules V and VI, along with the secondary motor representation in lobules VIIIa and VIIIb. Working memory activation convergence was limited to Lobule VI and Vermis VI; however, unlike in YA we did not see any convergence across studies in Crus I and II. With respect to long term memory, convergence across studies in OA appears to be more extensive than in YA, extending from Lobule VI to Crus II and also including Lobule IX. When looking at other cognitive tasks which were primarily those that tapped into executive functions, broadly defined, OA demonstrated significant overlap across studies in Crus I, Vermis VI, and Lobule V. Finally, for language tasks there was no significant convergence across studies in OA.

### Age Differences in Cerebellar Activation Overlap

Group differences in activation convergence across studies for all task domains, except for language, were computed (Figure 3, Table 7). Due to the nature of the ALE algorithm, comparisons cannot be made when one group does not show any significant activation across tasks. As such, we were unable to analyze language. With that said, it is worth noting that in YA, there was significant convergence across language tasks but this was not the case at all in OA, suggesting less reliable activation across studies in advanced age, perhaps due to less activation overall.

**Figure 3.**
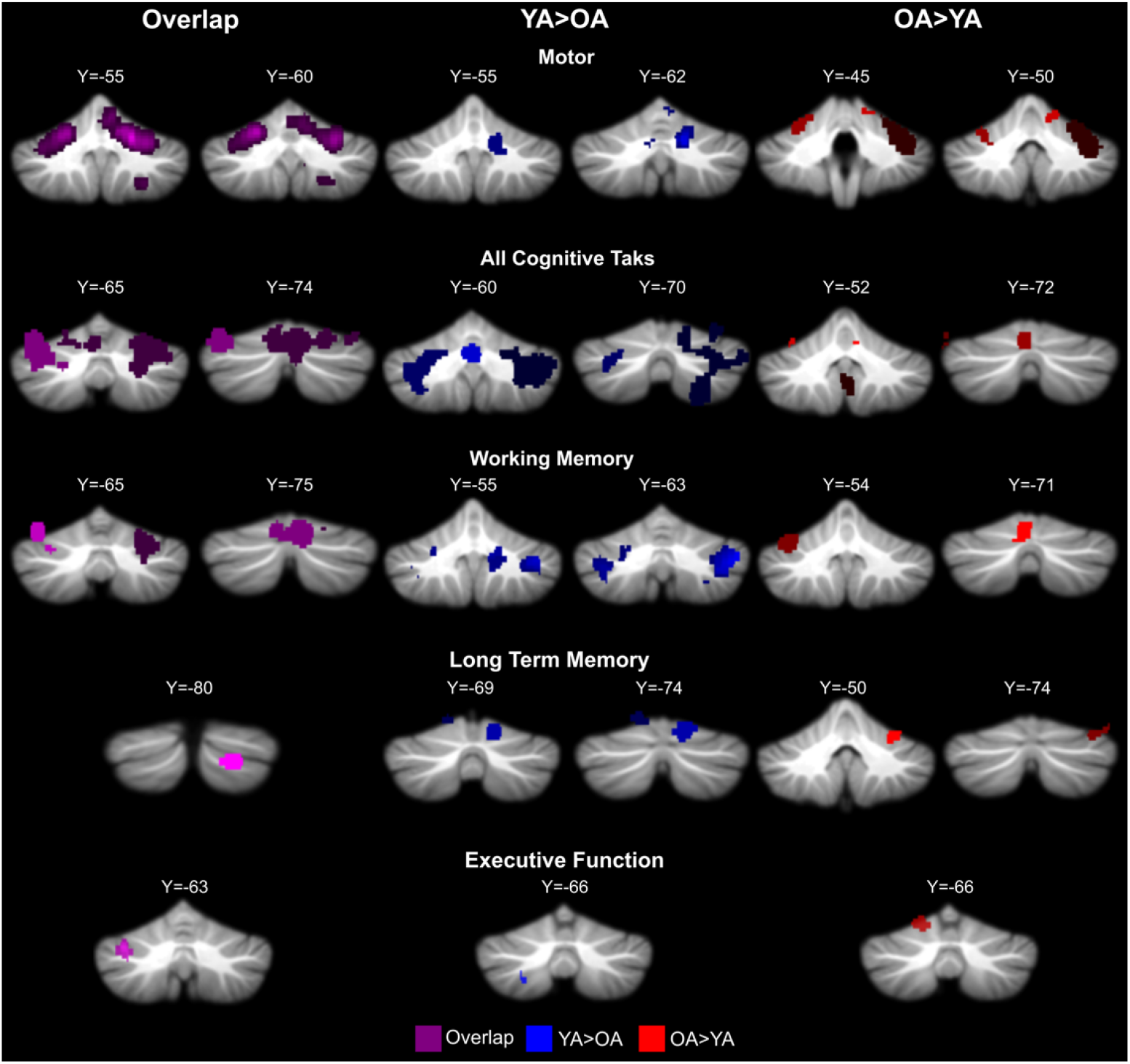
Overlap between age groups as well as age differences across studies in cerebellar activation. Because for several cognitive task domains there were not enough foci to compare the two age groups, or because there was no significant overlap within an age group, all of the cognitive task domains were combined and investigated together as well. Purple: overlap between age groups across tasks. Blue: YA>OA. Red: OA>YA.

**Table 7.**
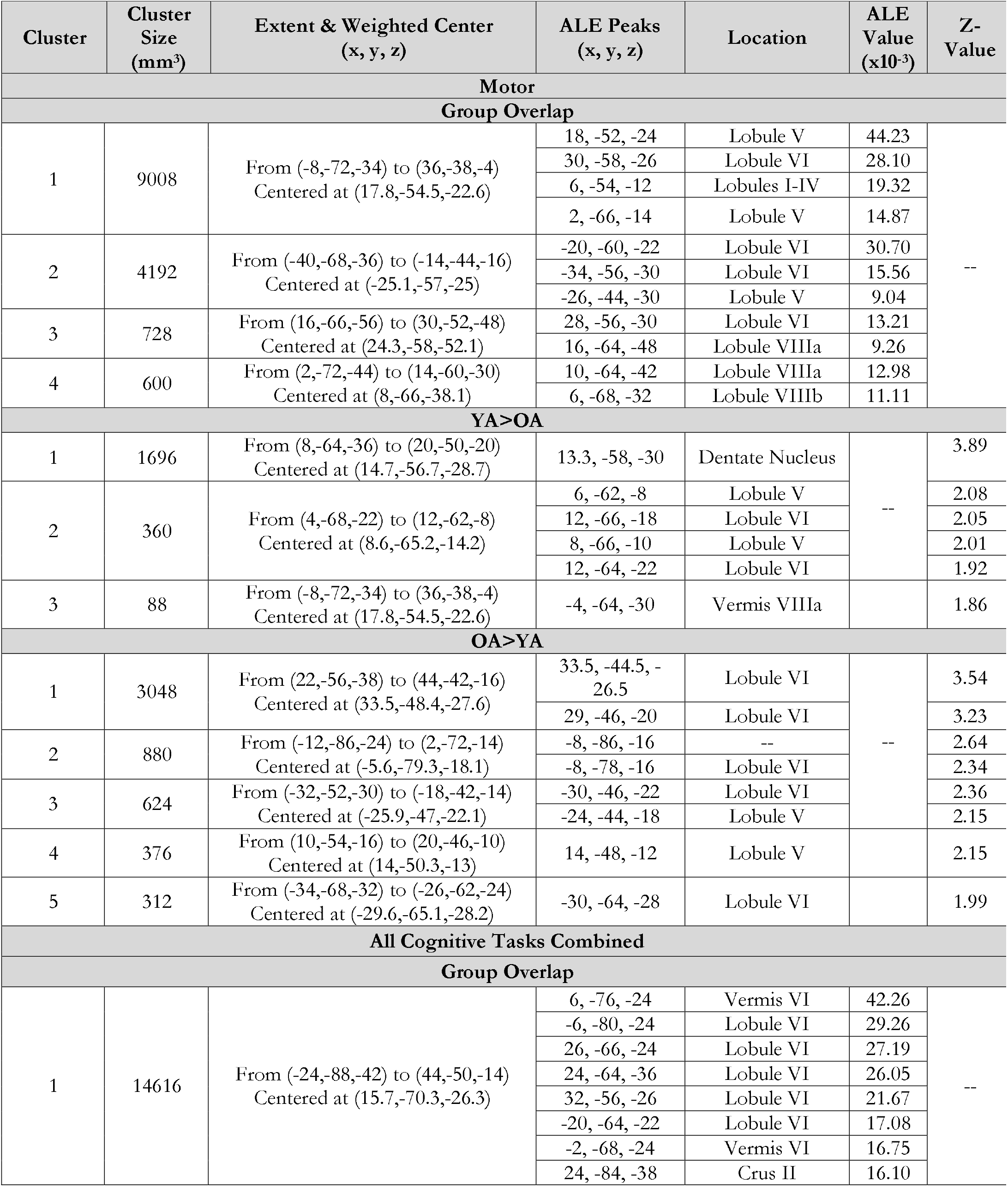
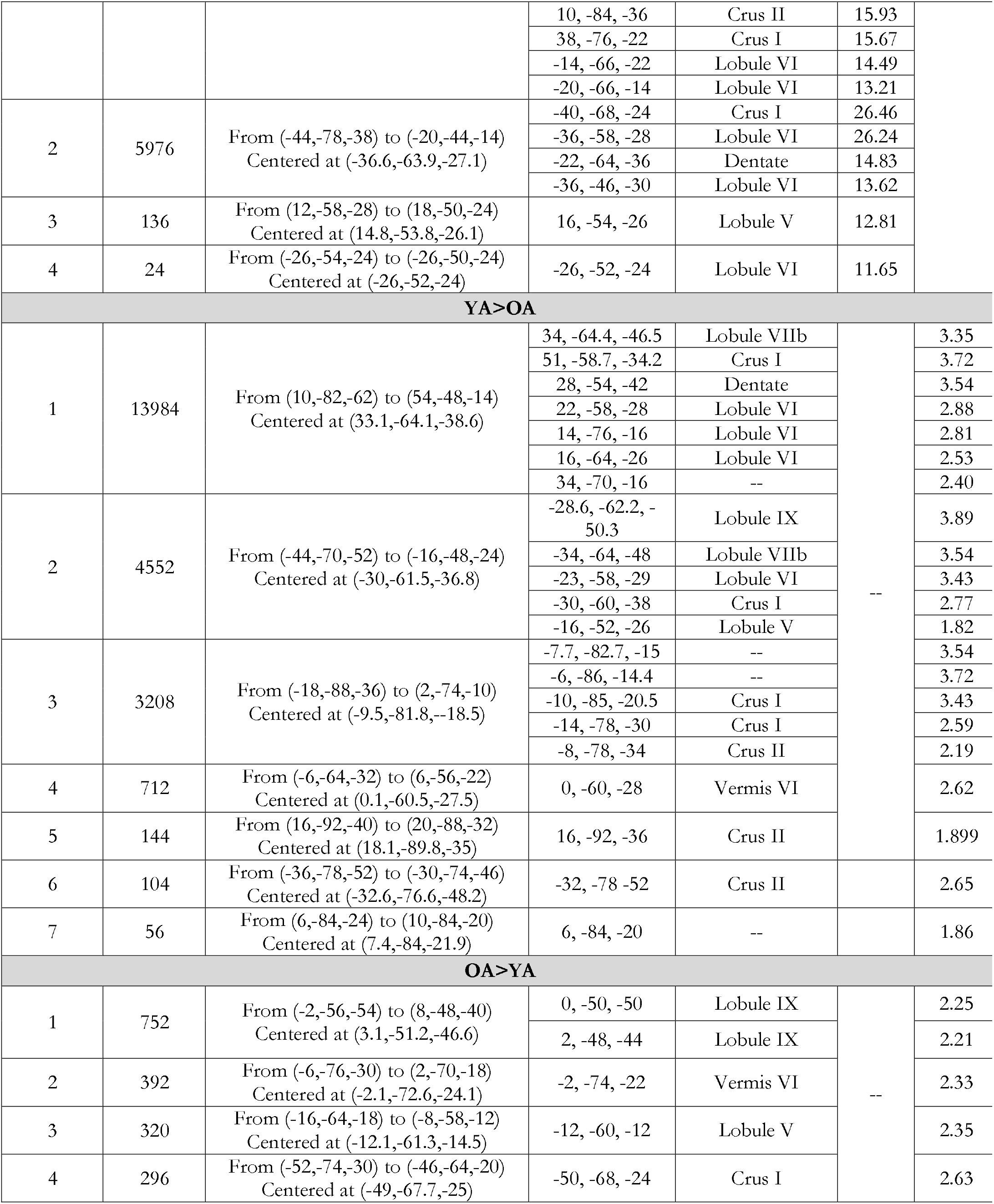
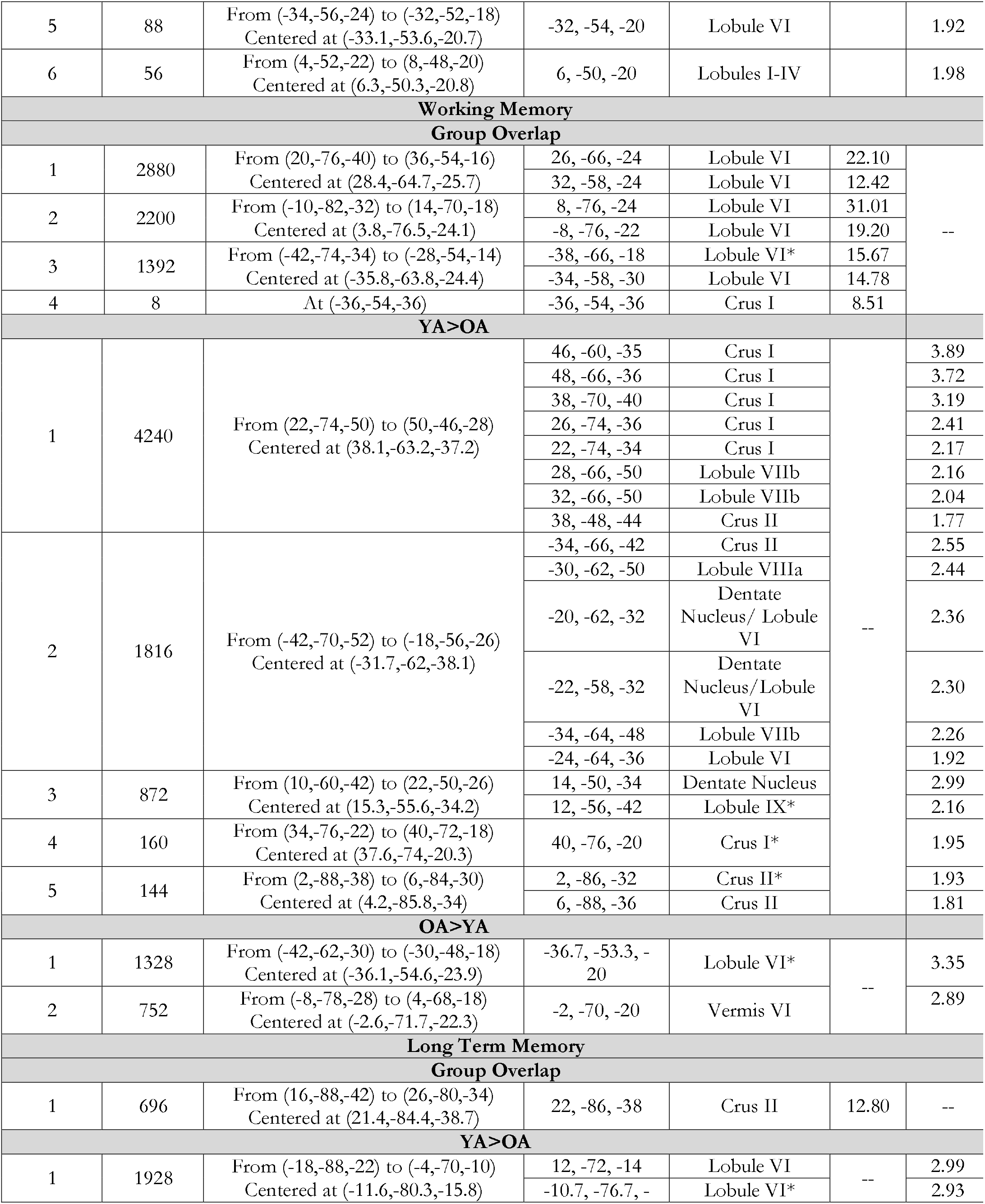
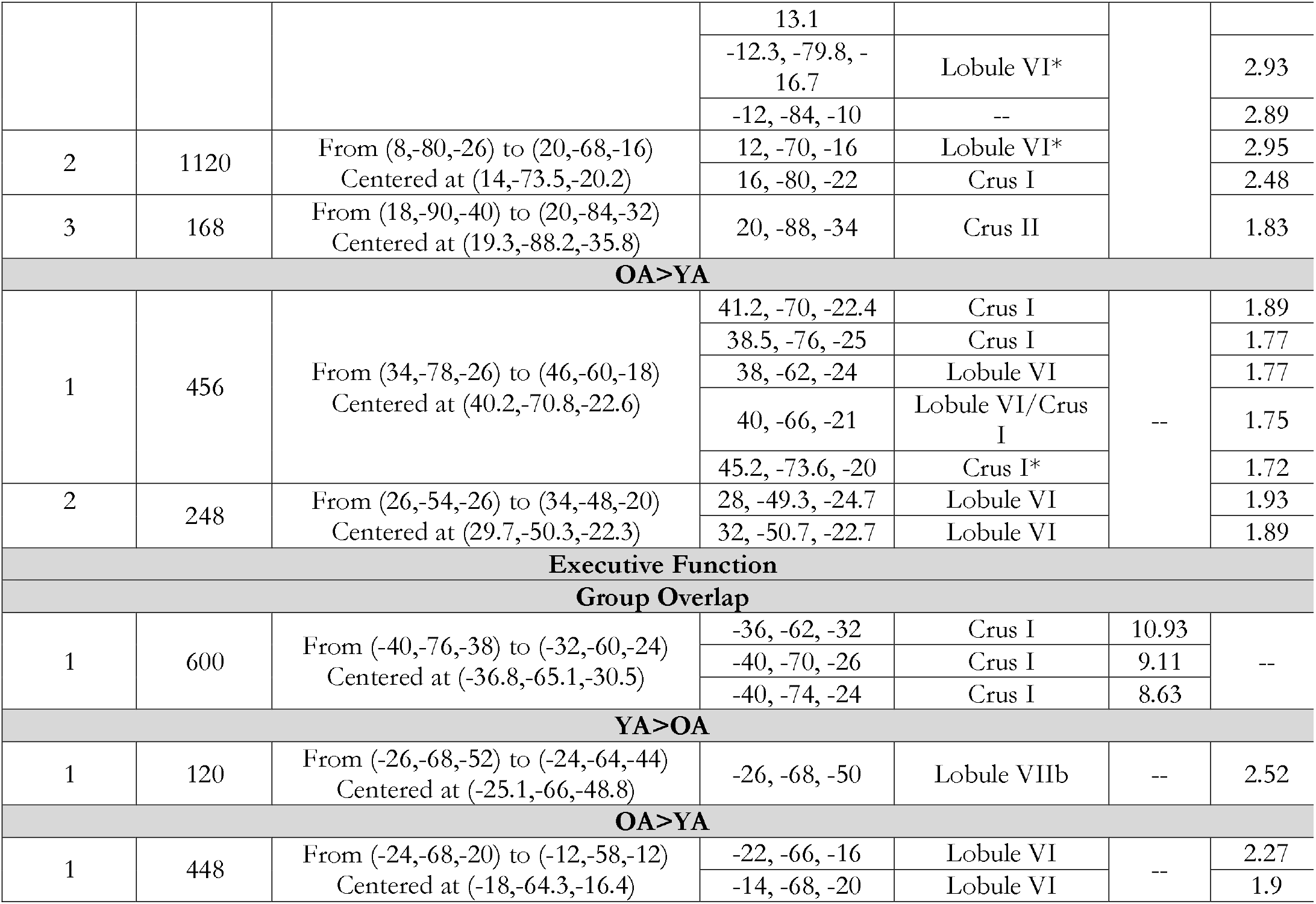
Group comparisons by task. *Peak outside of SUIT Atlas space, the closest region to the reported peak is listed.

With respect to motor tasks, it is first notable that there was significant overlap between the two age groups in regions of the anterior cerebellum, including Lobules I-IV, V and VI. YA showed greater convergence across studies in the dentate nucleus and Lobule V, an area heavily involved in motor processing (Stoodley & Schmahmann, 2009). In addition, some convergence extended into Lobule VI. In OA, convergence across studies relative to YA was seen primarily in Lobule VI. Notably, while greater convergence across studies in YA was limited primarily to the right hemisphere, in OA this was bilateral.

Working memory tasks also resulted in a great deal of activation overlap across studies when looking at the conjunction of the two age groups. Not surprisingly, this was localized to bilateral Lobule VI and left Crus I. Greater convergence across studies in YA was seen in Crus I and II, the dentate nucleus, Lobule VIIb, and Lobule VIIIa. Greater activation convergence in OA was much more limited and seen only left Lobule VI and Vermis VI. The spatial extent of the overlap unique to OA was just more than one quarter (28.7%) of that which was unique to YA. With respect to long term memory, there was some shared convergence across studies in both age groups localized to Crus II. Greater convergence in YA as compared to OA was seen in Lobule VI, Crus I, and Crus II. Similar lobules were observed when looking at areas where OA had greater convergence, but localization within these lobules was unique relative to YA, and again, the spatial extent of the convergence areas that were greater in YA was much larger. In this instance, the area in OA was only 21.9% of that seen in YA. In both of these memory domains this suggests that across studies in YA, there is more consistent activation across larger aspects of the cerebellum as compared to OA, where convergence was more limited in its spatial extent.

When investigating other cognitive tasks, which primarily includes executive function tasks, convergence in activation across studies was seen in Crus I and Vermis VI. When looking at the two groups relative to one another greater convergence in YA was seen in VIIIb, and greater convergence in OA was seen in Lobule VI. However, in both cases these were relatively small areas. Because there was no significant convergence across language tasks in OA, we were unable to conduct a group comparison.

Finally, we combined all cognitive tasks to compare overlap between YA and OA. This allowed us to include language in a broader analysis of group differences in convergence across tasks. Consistent with the broader literature suggesting that the lateral posterior cerebellum is involved in cognitive task processing and has connections (both structural and functional) with the prefrontal cortex (Bernard et al., 2012; Chen & Desmond, 2005; Krienen & Buckner, 2009; Salmi, Pallesen, & Neuvonen, 2010; Stoodley & Schmahmann, 2009), there was substantial convergence overlap between the age groups in hemispheric Crus I and Lobule VI, though Vermis VI was also implicated along with the cerebellar dentate nucleus. In YA, greater convergence was seen in a wide swath of the posterior cerebellum. This included Lobule VI, Crus I, and Crus II, as well as Lobule VIIb and Lobule I. In OA relative to YA, there was convergence in Lobule IX, Crus I, and Lobule VI, though there was also an area in Lobule V. Convergence in Lobules I-IV and V was unique to the OA sample. Most notably, the overall volume of the areas of increased overlap was substantially smaller in the OA group (1,634 mm^3^) as compared to the YA (22,760 mm^3^).

## Discussion

Here, using ALE meta-analysis, we directly compared cerebellar activation convergence across task domains in OA and YA for the first time. Our results indicate that YA and OA recruit cerebellar resources differently during task performance, as evidenced by group differences in areas of activation convergence across studies. However, there is also substantial overlap in the convergence patterns when comparting the two age groups. These findings represent several important practical and theoretical advances in our understanding of cerebellar contributions to behavior in advanced age. First, this expands our understanding of the cerebellum in aging beyond the anatomical and connectivity domains to include functional activation patterns. Second, more broadly, this extends our understanding of the neural underpinnings of task performance in OA to include the cerebellum. Though as this investigation demonstrates, cerebellar activation has long been found in functional imaging studies of aging; but, it has not been the focus of study. Concatenation across investigations in this manner provides a powerful tool to better understand cerebellar functional activation patterns in advanced age. Most notably, there is evidence to suggest potential under-recruitment of the cerebellum in individual cognitive domains, and when all cognitive tasks are investigated together. This is consistent with our predictions based on degraded connectivity and volume in the cerebellum in OA (Bernard & Seidler, 2014). However, the activation convergence patterns differ for motor tasks. There is less convergence in OA in the primary motor regions of the cerebellum relative to YA, but across studies, there is greater convergence in secondary cerebellar motor regions. Together, these results suggest that with advanced age, cerebellar resources are not relied upon as effectively and efficiently in OA during task performance.

Unlike in the cerebral cortex where an increase in bilateral activation and compensatory recruitment during cognitive task performance in OA has been reported (Cabeza, 2002; Cappell, Gmeindl, & Reuter-Lorenz, 2010; Reuter-Lorenz et al., 1999), here we demonstrate a relative decrease in convergence across studies investigating cognitive task domains. Though this does not directly indicate activation, it does imply that the organization of activation across studies is not consistent with cortical bilateral patterns of activation. We suggest that OA may not be consistently engaging bilateral regions of the cerebellum during cognitive task performance as they do in the cortex. Though surprising in the context of the cortical literature, this is consistent with the hypothesis put forth in our recent review (Bernard & Seidler, 2014). Because connectivity is lower in OA relative to YA, information exchange between the cortex and cerebellum may be degraded. As such, OA may not be able to effectively recruit cerebellar resources for information processing (Bernard & Seidler, 2014). The results here are consistent with this idea, and we suggest that this difference in the recruitment of the cerebellum may be particularly important for behavior. Specifically, cerebellar resources may be especially important scaffolding for performance in advanced age (e.g., Reuter-Lorenz & Park, 2014). Reliance upon more automatic processing in the cerebellum via internal models of behavior (Ramnani, 2014) would free up cortical resources and help maintain performance. However, those resources are not recruited consistently in OA as evidenced by the activation convergences patterns across studies seen here. Further, it may in fact be the case that the inability to utilize these cerebellar resources contributes, at least in part, to the bilateral cortical activation patterns seen in OA.

Somewhat surprisingly, we found a distinct pattern of convergence differences for motor tasks. OA showed significantly more convergence in Lobule VI compared to YA. This region has connectivity patterns with prefrontal cortical and premotor regions (e.g., Bernard et al., 2012; Krienen & Buckner, 2009) and also shows activation during cognitive task performance (Stoodley & Schmahmann, 2009). This convergence pattern is more consistent with activation patterns seen in the cortex in OA (e.g., Cabeza, 2002; Cappell et al., 2010; Reuter-Lorenz et al., 1999). Again, though not directly indexing activation, seeing a consistency in this pattern across studies suggests that perhaps OA are engaging these other cerebellar regions in a compensatory manner. Notably, recent work using a predictive motor timing task demonstrated increased activation in the lateral posterior cerebellum with increasing age (Filip et al., 2019). The authors suggest that this activation pattern may provide scaffolding for performance in OA (Filip et al., 2019), consistent with extant models of aging (e.g., Reuter-Lorenz and Park, 2014). However, given that we did not see this pattern for cognitive tasks, why such scaffolding is present for motor tasks raises interesting mechanistic and theoretical questions.

As described above, in our past work, we had hypothesized that lower connectivity between the cerebellum and cortex coupled with volumetric differences in OA relative to YA would result in decreased activation during task performance, indicative of an inability to rely upon cerebellar resources for performance (Bernard & Seidler, 2014). Data from studies administering cognitive tasks are consistent with this notion, while data from motor tasks better parallel cortical findings (e.g., Reuter-Lorenz & Cappell, 2008) and are consistent with recent work suggesting that the cerebellum provides scaffolding for motor performance in advanced age (Filip et al., 2019). Though seemingly contradictory, these findings may in fact be quite consistent with inputs to the cerebellum, particularly in the context of control theory (Ramnani, 2006). In this context, and as we previously proposed (Bernard & Seidler 2014), inputs and outputs between the cerebellum and cortex are degraded in advanced age, resulting in less efficient internal models, evidenced through decreased functional activation. However, a key part of these models is the function of the comparator, which compares the predicted behavior to its consequences (Ramnani, 2006). The inferior olive is suggested to be the comparator for both motor and cognitive processes, however the *input* to this region differs. For cognitive processes, input to the inferior olive comes from cortico-olivo-cerebellar pathways, while those for motor come from the spino-olivo-cerebellar pathways (Ramnani, 2006). We speculate that the spinal pathways are relatively intact, particularly as compared to the cortical pathways, and as such, OA are better able to recruit cerebellar resources during motor task performance. Thus, in the context of control theory, while the cerebellum may be capable of providing compensatory activation, because of the input from the cortex for the updating of internal models, these resources cannot be brought online effectively. Together, our findings suggest that in OA cerebellar resources may be under-recruited during cognitive tasks as evidenced by the relative decrease in convergence across studies when compared with YAs, and we propose that compensatory scaffolding during motor performance is due to spino-olivo-cerebellar inputs.

While the meta-analytic approach employed here allows for insights into cerebellar activation patterns across studies, it is not without limitations. Most notably, we were unable to account for behavioral performance and brain-behavior relationships. While an understanding of brain-behavior relationships is a key question moving forward, the inclusion of a behavioral meta-analysis here is beyond the scope of our work. Furthermore, there are numerous existing behavioral meta-analyses across domains demonstrating age differences in performance (e.g., Maldonado et al., In press; Verhaeghen et al., 2003; Wasylyshyn et al., 2011), negating the need for an additional behavioral meta-analysis here. Additionally, we did not conduct a complete search of the entire YA imaging literature. This would have resulted in large differences in statistical power between groups and potentially biased results in favor of the YA sample. As such, this is not a comprehensive investigation of cerebellar activation patterns in YA, but several meta-analyses on this topic have already been conducted (Stoodley & Schmahamann, 2009; E et al., 2012). Further, it is notable that even with less power in the YA data sample, we nonetheless see less overlap across studies in OA during cognitive task performance. If anything, this sample was biased in favor of seeing more consistent activation overlap in OA given the size of the sample; however, for cognitive task processing, the opposite pattern was demonstrated, providing powerful evidence for differences in cerebellar engagement with age. Finally, one of the greatest benefits of meta-analyses is the ability to concatenate across large literatures. However, this also means concatenating across studies with different methodological approaches, and varying degrees of information related to the study samples. As such, we included both PET and fMRI studies, consistent with past meta-analyses of cerebellar function (Stoodley & Schmahmann, 2009; E et al., 2012), as well as different inclusion and exclusion criteria. Notably however, the ALE algorithm accounts for uncertainty and variability across subjects and sites so as to be relatively robust to these methodological differences (REikhoff et al., 2012).s While this means we cannot carefully control for these individual factors, this work also provides a powerful indicator of activation patterns seen in different age groups, and the diversity in the samples is likely more representative of the broader population as a whole.

Together, this work represents the first comprehensive investigation into cerebellar activation patterns in OA. First, we demonstrated that during the performance of cognitive tasks, OA show *less* convergence in cerebellar foci across studies than YA, perhaps indicative of decreases in activation, in contrast to what is seen in the prefrontal cortex (e.g., Reuter-Lorenz et al., 1999; Cabeza, 2002; Reuter-Lorenz & Cappell, 2008), consistent with our hypothesis and prior work (Bernard & Seidler, 2014). We suggest that cerebellar processing is critical for optimal and efficient behavior. In advanced age, these resources are not brought online, likely due to degraded communication with the cortex (Bernard & Seidler, 2014). As such, OA are unable to use this critical region and scaffolding for performance, resulting in behavioral declines. Conversely, we see more extensive convergence across studies in OA during motor tasks. However, we propose that this is due to spinal afferents that bypass the cortex and allow for compensatory activation (Filip et al., 2019). Thus, on the basis of these findings we suggest that cerebellar functional activation differences with advanced age result in dissociable behavioral impacts due to the source of inputs through the inferior olive. While the cerebellum may be able to engage in compensatory activation for motor tasks, this is not the case in the cognitive domain.

